# Energetic coupling of an active site residue in penicillin-binding protein 2 from *Neisseria gonorrhoeae* with a resistance-associated conformational switch in the β3-β4 loop

**DOI:** 10.64898/2026.08.07.743578

**Authors:** Caleb M. Stratton, Sandeepchowdary Bala, Marissa M. Bivins, Robert A. Nicholas, Christopher Davies

**Affiliations:** University of South Alabama, Department of Biochemistry and Molecular Biology, Mobile, Alabama, 36688, United States; University of North Carolina at Chapel Hill, Department of Pharmacology, Chapel Hill, North Carolina, 27599, United States

## Abstract

Mosaic *penA* alleles encoding highly mutated variants of penicillin-binding protein 2 (PBP2) are the principal determinants of ceftriaxone resistance in *Neisseria gonorrhoeae*. Resistance-associated mutations in PBP2 from the ceftriaxone-resistant strain H041 restrict formation of the inward conformation of the β3–β4 loop associated with efficient acylation, but how β-lactam recognition is coupled to this conformational switch is unknown. Because the conserved active-site residue Tyr422 interacts with the R_1_ substituent of β-lactams, we investigated its role in coupling ligand recognition and acylation activity. Mutation of Tyr422 to Ala lowered acylation rates by up to 120-fold for cefoperazone and piperacillin, whereas acylation rates of ceftriaxone increased 4-fold. Unexpectedly, the crystal structure of the Y422A mutant acylated by ceftriaxone revealed that the β3–β4 loop had adopted the inward, high-activity conformation, despite position 422 being spatially distant from the loop. Transformation experiments showed that cell viability requires a tyrosine at position 422, indicating the residue is essential for transpeptidase function. Together, these findings reveal an energetic coupling between an active-site residue in PBP2 and a conformational switch whose equilibrium is altered by resistance mutations. The previously observed higher activity of β-lactams containing extended R_1_ groups is consistent with stronger interactions with Tyr422 that favor the conformational switch. Molecular modeling suggests that such groups enhance activity by mimicking the *iso*-Glu region of the pentapeptide substrate. Overall, we propose that access to the high-activity state of PBP2 where the β3–β4 loop is inward is regulated by interactions between Tyr422 and β-lactam R_1_ groups, and that resistance mutations function by tilting the balance toward a lower activity state.

## Introduction

The Gram-negative diplococcus *Neisseria gonorrhoeae*, the etiological agent for the sexually transmitted infection (STI) gonorrhea, is an international risk to public health. In the United States, gonorrhea is the second most reported bacterial STI, with approximately 600,000 new cases reported in 2020 and 82 million infections per year worldwide ^3^. Untreated gonorrhea can have serious health consequences, including infertility due to prolonged inflammation of the reproductive tract, disseminated gonococcal infection (DGI), and an increased risk of both contracting and transmitting HIV ^4, 5^.

The public health risk from *N. gonorrhoeae* is growing due to the emergence of strains exhibiting resistance to the extended-spectrum cephalosporin (ESC) ceftriaxone, the current first-line treatment and the only antibiotic recommended by the US Centers for Disease Control (CDC) for gonorrhea ^6–13^. For example, the first ESC-resistant strain to be isolated (H041) exhibits a 2,000-fold higher minimum inhibitory concentration (MIC) for ceftriaxone compared to the penicillin– and cephalosporin-susceptible strain FA19 ^10, 14, 15^. The key determinant for resistance in ESC-resistant strains is the presence of mosaic *penA* alleles that encode highly mutated variants of penicillin-binding protein 2 (PBP2), as demonstrated by elevation of the MIC to levels above the clinical breakpoint when the *penA* allele from H041 (*penA41*) was transformed into FA19 ^15^. Consistent with this, purified PBP2 derived from H041 (PBP2^H041^) exhibits a 2,000-fold lower second-order rate of acylation for ceftriaxone compared to PBP2 from FA19 (PBP2^FA19^) ^2, 15^.

The sequences of PBP2^FA19^) and PBP2^H041^ differ by 61 amino acids. Of these, a subset of seven contributes the majority of resistance to ESCs ^16^. Most of the key mutations are in the active site region, including two (F504L and N512Y) on the β3-β4 loop. Crystal structures show the loop to occupy at least two states: an inward position near the active site that correlates with higher acylation activity and a so-called extended “outbent” conformation away from the active site that correlates with lower acylation activity^2, 17^ (Fig. 1). The F504L and N512Y mutations appear to restrict inward movement of the loop, thereby hindering formation of the high-activity state ^2, 17^.

**Fig. 1:**
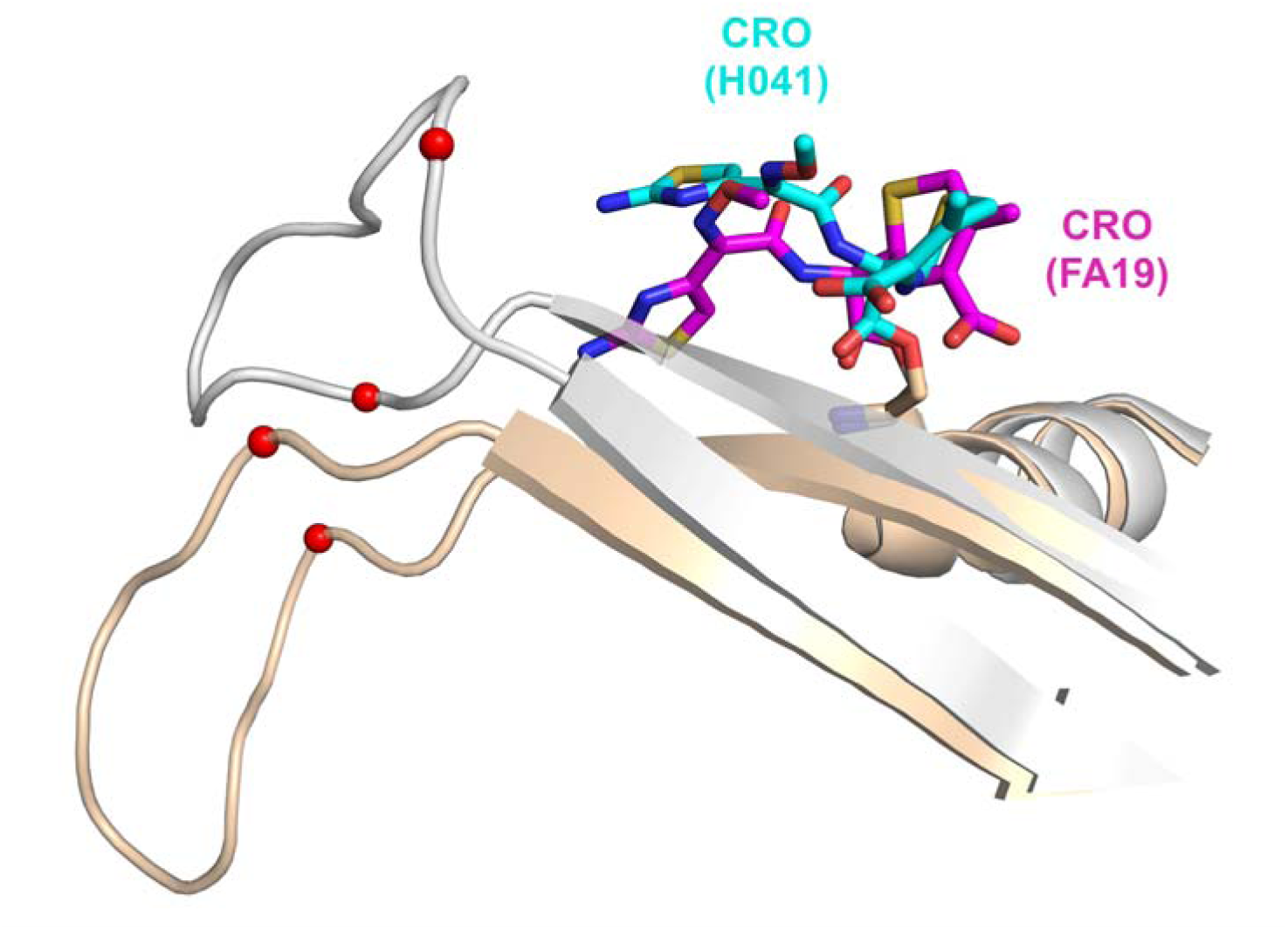
The β3-β4 loop occupies different conformations in *N. gonorrhoeae* PBP2 from the ESC-susceptible strain FA19 and the ESC-resistant strain H041. PBP2 from FA19 is colored grey with ceftriaxone (CRO) in magenta bonds and PBP2 from H041 is colored wheat and ceftriaxone is cyan. The locations of the resistance mutations are marked with

In its inward position, residues of the β3-β4 loop, including Arg502, Glu307 and Tyr509, form a cluster of interactions centered around the β-lactam R_1_ group. By forming a stacking interaction with the R_1_ group, Tyr422, which emanates from the α8-α9 loop on the opposite side of the active site from β3, is also involved in this clustering. Notably, the relative positions of the tyrosyl side chain and the aminothiazole R_1_ group of ceftriaxone differ between acylated structures of PBP2^FA19^ and PBP2^H041^. In PBP2^H041^, the aminothiazole group stacks outside the phenolic side chain of Tyr422 relative to the surface of the protein, whereas in PBP2^FA19^, it is underneath Tyr422 (Fig. 2). Furthermore, the outside position of the R_1_ group in PBP2^H041^ correlates with the outbent position of the β3-β4 loop, and the inside position in PBP2^FA19^ correlates with the β3-β4 loop being inward. However, it is unclear whether this is a cause-and-effect relationship and indeed whether the different positions of Tyr422 relative to the aminothiazole impact acylation activity.

**Fig. 2:**
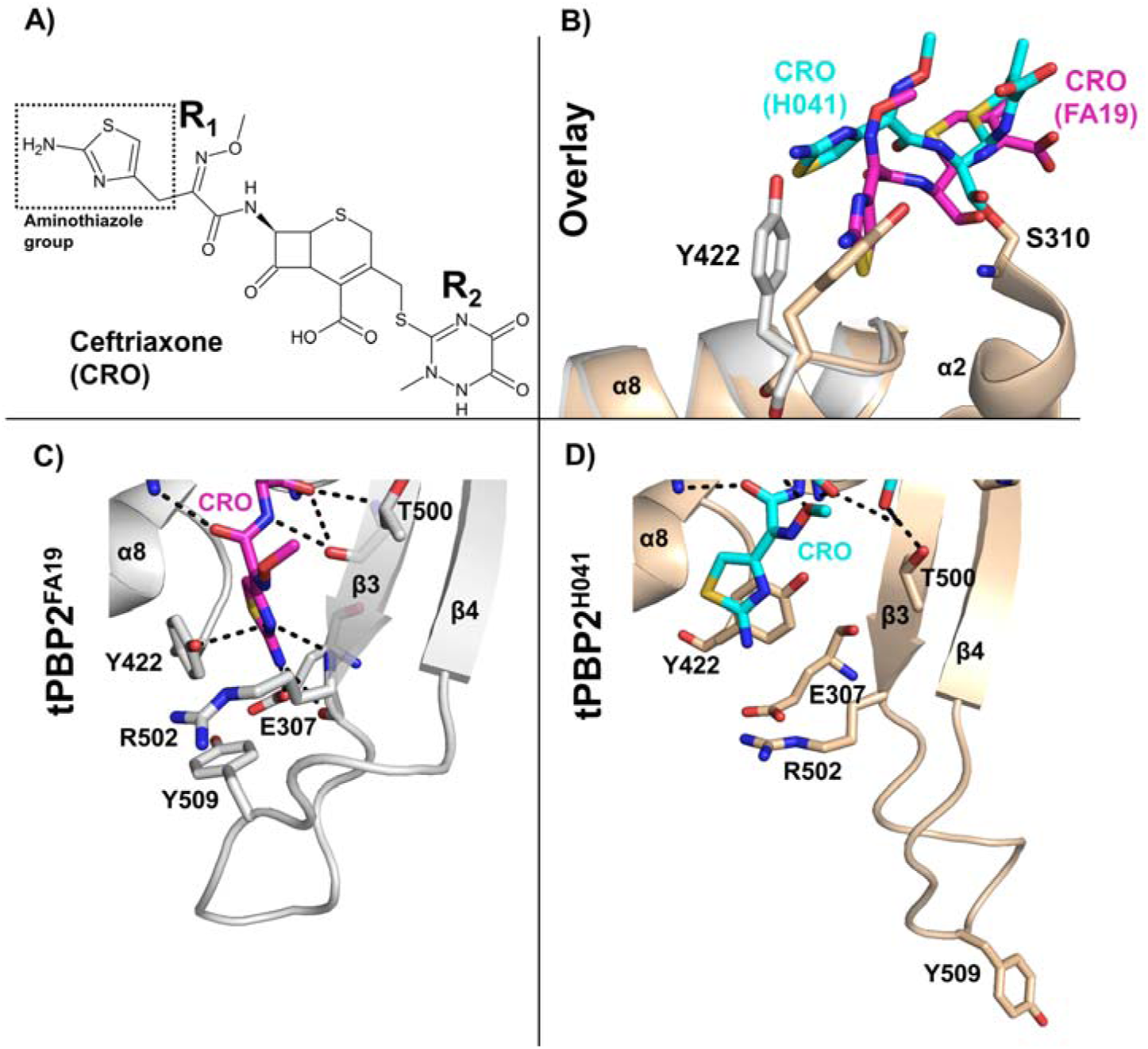
The R_1_ aminothiazole group of ceftriaxone interacts differently with Tyr422 in PBP2^FA19^ compared to PBP2^H041^. **A)** Molecular structure of ceftriaxone (CRO). **B)** Superposition of the ceftriaxone-acylated structures of tPBP2^FA19^-CRO (grey/magenta) superimposed onto tPBP2^H041^ (wheat/cyan). **C)** Active site of tPBP2^FA19^ acylated by ceftriaxone ^1^, where ceftriaxone is colored with magenta bonds. **D)** Active site of tPBP2^H041^ acylated by ceftriaxone ^2^, where ceftriaxone is colored with orange bonds. Potential hydrogen bonds depicted as dashed lines.

Recently, we reported the acylation rates for several β-lactams against PBP2^H041^, including a panel of cephalosporins and ureidopenicillins. The resulting structure-activity relationship (SAR) indicated that the R_1_ group plays a crucial role in the activity of β-lactams ^18^ and that higher acylation activity correlates with the β3-β4 loop occupying the inward position ^19^. Here, we examine whether the interaction between the R_1_ group and Tyr422 affects acylation rates for β-lactams and whether this alters the behavior of the β3-β4 loop. We find that acylation rates of cefoperazone and piperacillin are significantly reduced against a Y422A mutant of PBP2^H041^ compared to unmutated PBP2^H041^, whereas acylation rates for ceftriaxone increase. A crystal structure of PBP2^H041^-Y422A acylated by ceftriaxone shows that β3-β4 loop occupies the inward position instead of the outbent conformation when the protein is unmutated. These findings suggest that the tyrosine side chain acts as a “speed bump” against acylation by ceftriaxone but enhances inhibition by piperacillin and cefoperazone. They also reveal an unexpected allosteric connection between Tyr422 and the β3-β4 loop, even though these are spatially separate in the protein, and how this connection is exploited by resistance mutations.

## Materials & Methods

*Materials.* Ceftriaxone, cefixime and cefoperazone were purchased from Thermo Scientific Chemicals, Bocillin-FL and piperacillin were from Sigma Aldrich (St. Louis, MO), and ceftizoxime was from MedChemExpress (Monmouth Junction, NJ).

*Protein expression and purification.* The cloning, expression and purification of a truncated construct of PBP2^H041^ (tPBP2^H041^) comprising the transpeptidase domain of the protein has been described previously ^1^. This construct expresses the TPase domain as a fusion with maltose-binding protein (MBP) with an intervening site for cleavage by TEV protease and a His_6_ at the N terminus. Constructs are denoted as ‘tPBP2*^allele^*-X’, where ‘t’ signifies the truncated transpeptidase domain, the superscript identifies the strain of *N. gonorrhoeae* from which the *penA* allele is derived, and ‘X’ represents a site-directed mutation. The truncated constructs are acylated by Bocillin-FL at the same rate as their full-length protein equivalents ^20^. Y422A and Y422F mutations were introduced into the tPBP2^H041^ construct by a commercial service (GenScript) and the resulting plasmids transformed into BL21 (DE3) competent cells (Sigma Aldrich). Mutant proteins were purified as reported for tPBP2^H041 2^. In brief, after expression in *E. coli* BL21 (DE3) cells and cell lysis, fusion proteins were purified by nickel-affinity chromatography, digested with His_6_-tagged TEV protease, and a second nickel-affinity column was used to separate tPBP2 from MBP, His_6_-TEV protease and undigested fusion protein. As a final step, purified proteins were buffer-exchanged into Tris-HCl buffer, pH 7.8, with 10% glycerol and 500 mM NaCl, and concentrated to ∼13 mg/mL, and frozen aliquots were stored at –80°. Typical purities are ∼95%, as judged by SDS-PAGE.

*Determination of second-order acylation rate constants*. Second-order rate constants for the tPBP2^H041^-Y422A and Y422F mutants were determined using a time-course kinetic experiment ^18, 19^. Each protein (1 µM) was incubated at room temperature with Bocillin-FL ^21^ (5, 6, 8, 10 and 12 µM) in 50 mM phosphate-buffered saline (PBS, pH 8.0) for 0 to 3,600 seconds. At set intervals, 20 µL aliquots were withdrawn and mixed with 5 µL of 4X non-reducing SDS loading buffer to quench the reaction (Thermo Scientific Chemicals). Samples were heated to 95°C for five minutes and electrophoresed on 12% Mini-Protean TGX SDS-PAGE gels (Bio-Rad). Gels were imaged on a ChemiDoc^TM^ MP Imaging System to detect Bocillin-FL and then stained with Coomassie to measure protein levels. Densitometry was performed using ImageJ ^22^. The acylation rate of Bocillin-FL for each mutant was then calculated as described previously ^18^.

The acylation rates of β-lactam antibiotics for tPBP2^H041^-Y422A and Y422F were determined in a competition assay by measuring the concentration of β-lactam required to inhibit the binding of Bocillin-FL by 50% (IC_50_). Reactions were carried out with 1 µM of protein mixed simultaneously with 100 µM Bocillin-FL and increasing concentrations (1 µM to 10 mM) of antibiotic dissolved in 50 mM PBS at pH 8.0. The reactions were allowed to incubate for 1 hour at room temperature and then quenched with 5 µL of 4X non-reducing SDS loading buffer. Samples were heated to 95°C for 5 minutes prior to running on SDS-PAGE. Gels were subsequently imaged as described above. Second-order acylation rate constants (*k*_2_/K_S_) of the β-lactams were calculated as follows:

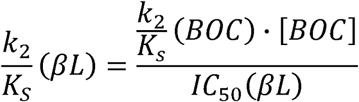

where *k*_2_/K_S_ (BOC) is the second-order acylation rate constant of Bocillin-FL, [BOC] is the concentration of Bocillin-FL used in the competition experiment, IC_50_ (βL) is the half-maximal inhibitory concentration of the β-lactam, and *k*_2_/K_S_ (βL) is the second-order acylation rate constant of the β-lactam ^23^. Values were obtained from a minimum of three separate experiments.

*Crystallization*. Both tPBP2^H041^-Y422A and tPBP2^H041^-Y422F were crystallized in the same conditions as previously described for tPBP2^H041 2^. Briefly, the proteins (13 mg/mL) were crystallized using a Gryphon liquid dispensing system (Art Robbins, Sunnyvale, CA, USA) in a 96-well sitting drop format, where 200 nL of protein was mixed with 200 nL of well solution. Microseeds of tPBP2^H041^ crystals were added to facilitate crystal growth. Crystals were obtained after 3-4 days incubation at 18°C over wells containing 0.1 M CHES buffer, pH 9.1 to 10.1, and 32-42% PEG 600.

*X-ray data collection and model refinement*. Crystals were flash-frozen in liquid nitrogen without adding additional cryo-protectant. Complexes with ceftriaxone were generated by addition of 1 µL of 10 mM ceftriaxone dissolved in 50 mM PBS added directly to the wells containing crystals, incubated for 5 min, and then flash-frozen in liquid nitrogen. For both mutants, diffraction data were collected at a wavelength of 1.00 Å on a Eiger-16M detector at the SER-CAT 22-ID beamline at the Advanced Photon Source in Argonne, IL, USA. 200° of data were collected in 0.25° oscillations with an exposure time of 0.2 s/frame and a crystal-to-plate distance of 230 mm. Data were processed using HKL2000 ^24^ and structures solved by refinement against *apo* tPBP2^H041^. Bound ligands were modeled into the |F_o_|-|F_c_| difference electron density map, followed by iterative cycles of model building and refinement using COOT ^25^ and PHENIX ^26^ or REFMAC ^27^. Ramachandran plots were calculated using MOLPROBITY ^28^.

*Thermal stability studies*. Thermal stability assays were conducted on a CFX Connect Real-Time PCR Detection System (Bio-Rad) using SYPRO Orange dye (Sigma), which fluoresces upon binding to hydrophobic protein regions exposed during denaturation (λex/λem = 470/570 nm). Reactions (25 µl) were run in triplicate with 10 µM protein in 10 mM Tris pH 8, 200 mM NaCl, and 5% glycerol. Protein was pre-incubated with β-lactam (10 µM, 50 µM, 100 µM, and 1 mM) for 30 minutes at room temperature before adding SYPRO Orange dye to a final 5X concentration (starting from the 5,000X stock from Sigma). Samples were heated from 10 to 90 °C at a rate of 1.0 °C/min. The resulting melting curves were analyzed in the CFX Maestro software (Bio-Rad), which calculates the negative derivative of relative fluorescence units (RFUs) at each temperature [-d(RFU)/dT]. The peak of this graph corresponds to the T*m* of the protein. T*m* values are reported as the mean ± the standard error of the mean (SEM) for three replicate experiments.

*Bioinformatic analysis.* A homology-derived secondary structure analysis (HSSP) ^29^ dataset was obtained by querying the MRS server ^30^ with the PDB code 6P53 (truncated tPBP2^FA19^). The HSSP databank retrieves aligned sequences based on the following formula:

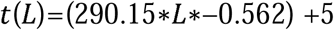

swhere L is the alignment length and t(L) is the percentage sequence identity. For a sequence to be included in the multiple-sequence alignment, the t(L) must be greater than or equal to a 55% threshold ^29, 31^. This resulted in 1,013 aligned sequences with predicted structural similarity. To diversify the set and minimize redundancy, only the first genus of each entry was retained, and subsequent entries from the same genus were removed. Unidentified sequences and those listed as “deleted” in UniProt were also excluded. This yielded a final dataset of 224 aligned sequences used to calculate the percent identity of amino acids at each position relative to the deposited structure. Interatomic contact surface areas for the aminothiazole group of ceftriaxone and the 2,3-dioxopiperazine ring of piperacillin/cefoperazone were calculated using the dr-sasa server ^32^. To assess the level of conservation of Tyr422 within *Neisseria* species, an alignment of all unique *penA* genes in the PubMLST database ^33^ was constructed using the Locus Explorer plugin.

*Generation of mutant PCR products.* Genomic DNA from FA19, FA19 *penA19-G545S*, and FA19 *penA41* strains was isolated using a Wizard Genomic DNA Purification Kit from Promega (Madison, WI) and used as template DNA for PCR amplifications. For the generation of mutant *N. gonorrhoeae* strains, four-primer PCR was used to create mutant *penA* genes. The primers included two overlapping mutagenic primers centered at the mutation site in opposite directions and two outside primers, with the forward primer starting at bp 135 of the *penA* gene and the reverse primer starting at bp 93 of the *murE* gene downstream of *penA* (Table S1). To increase efficiency of transformation, the 10-base pair gonococcal uptake sequence (5’-GCCGTCTGAA-3’) was appended to the 5’ end of the outside forward primer. For transformations using *penA* from FA19, a silent AccI restriction site was incorporated into the two mutagenic primers to help in screening for transformants. For each construct, the fragments from the two half reactions were cut from a 1% agarose gel and the gel pieces were spun through Xcluda D filter tips (Bio-Rad, Hercules, CA). The filtrates were then added to a new PCR tube and the full-length fragment was amplified with the outside primers. The PCR products were purified, quantified using a Nanodrop, and used for transformation into *N. gonorrhoeae*.

*Transformation of N. gonorrhoeae strains*. For transforming the *penA41-Y422X* mutant constructs into FA19, a standard liquid transformation protocol utilizing the capacity of *N. gonorrhoeae* to undergo natural recombination was used ^34^. Piliated colonies of *N. gonorrhoeae* strain FA19 were diluted to an OD_600_ of 0.18 in GC Medium Base (GCB) broth with Kellogg’s supplements I (4 mg/mL glucose, 0.1 mg/mL L-glutamate, 0.2 µg/mL co-carboxylase) and II (5 µg/mL Fe(NO_3_)_3_), 10 mM MgCl_2_, and 20 mM sodium bicarbonate. The PCR products were diluted into 1X SSC (0.15 M NaCl, 0.015 M sodium citrate) to ∼15 µg/mL in a final volume of 100 µL. The diluted FA19 culture (900 μL) was added to the DNA mixture and incubated at 37 °C, 5% CO_2_ for 5 hours, at which time 300 µL aliquots of the transformation mix were pelleted, resuspended in 50 µL of carry-over supernatant, and plated on GCB Agar plates containing 0.1 µg/mL cefixime as the selection agent. For transformations of Y422X mutants of *penA19-G545S*, selection was on 0.003-0.006 μg/ml cefixime. For transformation of *penA19* with only Y422X, there was no selection agent.

*Screening of transformants*. After overnight incubation, six colonies of each transformation were passaged and colony PCR of each transformant was conducted. Briefly, four to six colonies were mixed in a microcentrifuge tube with 15 µL 10 mM Tris pH 8.0, 1 mM EDTA, boiled for 5 minutes to lyse *N. gonorrhoeae* cells and centrifuged to pellet cell debris. The supernatants were used as DNA template for PCR with 5’ *penA_*45aa (without uptake sequence) and 3’ *murE*_31aa primers to amplify the *penA* gene. Successful PCR reactions were then purified and sent to sequencing to assess transformation of strains. For transformations of Y422X mutants of *penA19* and *penA19-G545S*, the PCR products were digested with AccI to identify those with the silent restriction site.

## Results

### Binding mode of the R1 group of piperacillin in tPBP2^FA19^

Previous crystal structures of the transpeptidase (TPase) domain of PBP2 from H041 (tPBP2^H041^) in complex with ceftriaxone show that the R_1_ aminothiazole moiety of ceftriaxone is on the inside of the sidechain of Tyr422 in tPBP2^FA19^ relative to the surface of the molecule, whereas in tPBP2^H041^, this moiety is on the outside^1, 2^ (Fig. 2). This suggests that the juxtaposition of the aminothiazole moiety and Tyr422 side chain is impacted by the resistance mutations present in tPBP2^H041^. We have also solved a crystal structure of tPBP2^H041^ in complex with piperacillin, which has an 85-fold higher *k_2_*/K_s_ acylation rate than ceftriaxone, and observed the 2,3-diketopiperazine R_1_ group to be outside of Tyr422 *i.e.,* in the same position relative to the R_1_ group in the ceftriaxone-tPBP2^H041^ complex ^19^. It was unknown, however, whether in the absence of resistance mutations, the 2,3-dioxopiperazine R_1_ group of piperacillin would be inside or outside of Tyr422 in PBP2^FA19^. To address this, we solved the crystal structure of PBP2^FA19^ acylated by piperacillin. The structure was determined at 3.15 Å from crystals in the same P2_1_ system as for the tPBP2^FA19^ construct, where there are two tPBP2^FA19^ molecules in the asymmetric unit (molecules A and B)(Table S2). Density for piperacillin is observed in both molecules of the active site, with stronger density seen in molecule A (Fig. 3).

**Fig. 3:**
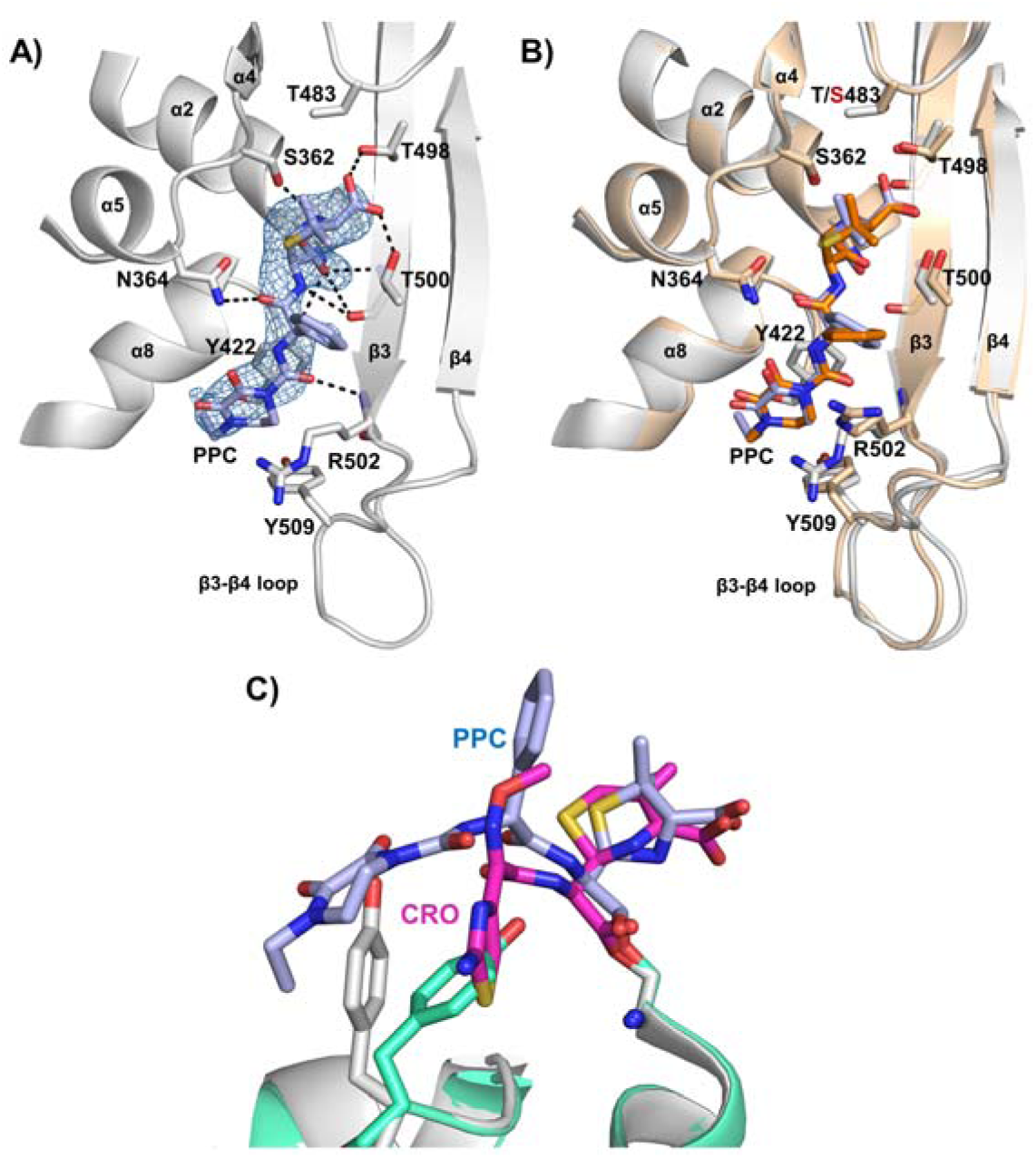
The structure of tPBP2^FA19^ acylated by piperacillin. **A)** Structure of tPBP2^FA19^ acylated by piperacillin (PPC), where the protein backbone is gray and PPC is shown with light blue bonds. Molecule A of the asymmetric unit is shown. The |Fo|-|Fc| difference electron density map is shown as a blue mesh and potential hydrogen bonds are shown as black dashed lines. **B)** Superimposition of the structures of tPBP2^FA19^ (gray backbone/light blue bonds) and tPBP2^H041^ (wheat backbone/orange bonds) acylated by piperacillin. **C)** Side view of tPBP2^FA19^ acylated by piperacillin (green backbone/light blue bonds) and by ceftriaxone (gray backbone/magenta bonds).

Comparison of the piperacillin-acylated structures of tPBP2^FA19^ and tPBP2^H041^ show them to be similar, with molecule A of tPBP2^FA19^ superimposing onto tPBP2^H041^ with an RMSD of 0.84 Å (Fig. 3B). Most pertinently, in contrast to the situation with ceftriaxone when bound to tPBP2^FA19^, the diketopiperazine ring of piperacillin is outside of Tyr422 (Fig. 3C), showing that unlike ceftriaxone, the position of the R_1_ group of piperacillin is not altered by resistance mutations. The interaction surface between the diketopiperazine ring and active site of tPBP2^FA19^ is 169 Å^2^, which is 26 Å^2^ larger than the interaction surface for the aminothiazole R_1_ of ceftriaxone inside of the Tyr422 binding pocket (143 Å^2^). An additional observation is that the β3-β4 loop occupies the same inward position as seen when tPBP2^H041^ is acylated by piperacillin ^19^. Hence, while the resistance mutations present in tPBP2^H041^ may affect the position of the aminothiazole group of ceftriaxone relative to the tyrosyl sidechain, that is not the case with the larger 2,3-dioxopiperazine moiety of piperacillin.

### Tyr422 decreases inhibition by ceftriaxone and cefixime but increases inhibition by ureido **β**-lactams

To determine whether the interaction between the piperacillin and ceftriaxone R_1_ groups and Tyr422 is responsible for the 85-fold difference in acylation rate against tPBP2^H041 19^, a Y422A mutation was introduced into tPBP2^H041^ (tPBP2^H041^-Y422A) and the second-order acylation rates for several β-lactams were determined using an SDS-PAGE gel-based competition assay with the fluorescent penicillin, Bocillin-FL ^18, 21^ (Fig. S1).

The acylation rates of cefoperazone and piperacillin were markedly lower for tPBP2^H041^-Y422A compared to those reported for tPBP2^H041 19^. Specifically, the acylation rates of cefoperazone and piperacillin decreased 4-fold and 49-fold, respectively, whereas the rates for ceftriaxone and cefixime increased 4– and 2-fold, respectively (Table 1). The slight difference between ceftriaxone and cefixime may be a consequence of ceftriaxone possessing a C3 leaving group, whereas cefixime does not. To examine whether the altered rates were due to the loss of packing against the tyrosyl ring of Tyr422 or the hydroxyl group, we also made a Y422F mutant. Rates for cefoperazone against this mutant were unchanged, whereas the rate for piperacillin increased to the point where it could not be measured accurately in the gel-based assay (Table 1). In contrast, the rates for ceftriaxone and cefixime decreased 10– and 34-fold, respectively. Taken together, these results suggest that Tyr422 is important for acylation of PBP2 by β-lactams, but has a differential effect depending on the β-lactam. Specifically, Tyr422 in tPBP2^H041^ enhances the acylation rate of β-lactams with the larger 2,3-dioxopiperazine ring, *i.e.,* cefoperazone and piperacillin, but lowers the acylation rate for those with smaller aminothiazole groups, *i.e.,* ceftriaxone and cefixime. In addition, the tyrosyl hydroxyl is more critical for β-lactams with smaller R_1_ groups than those with larger groups.

**Table 1:**
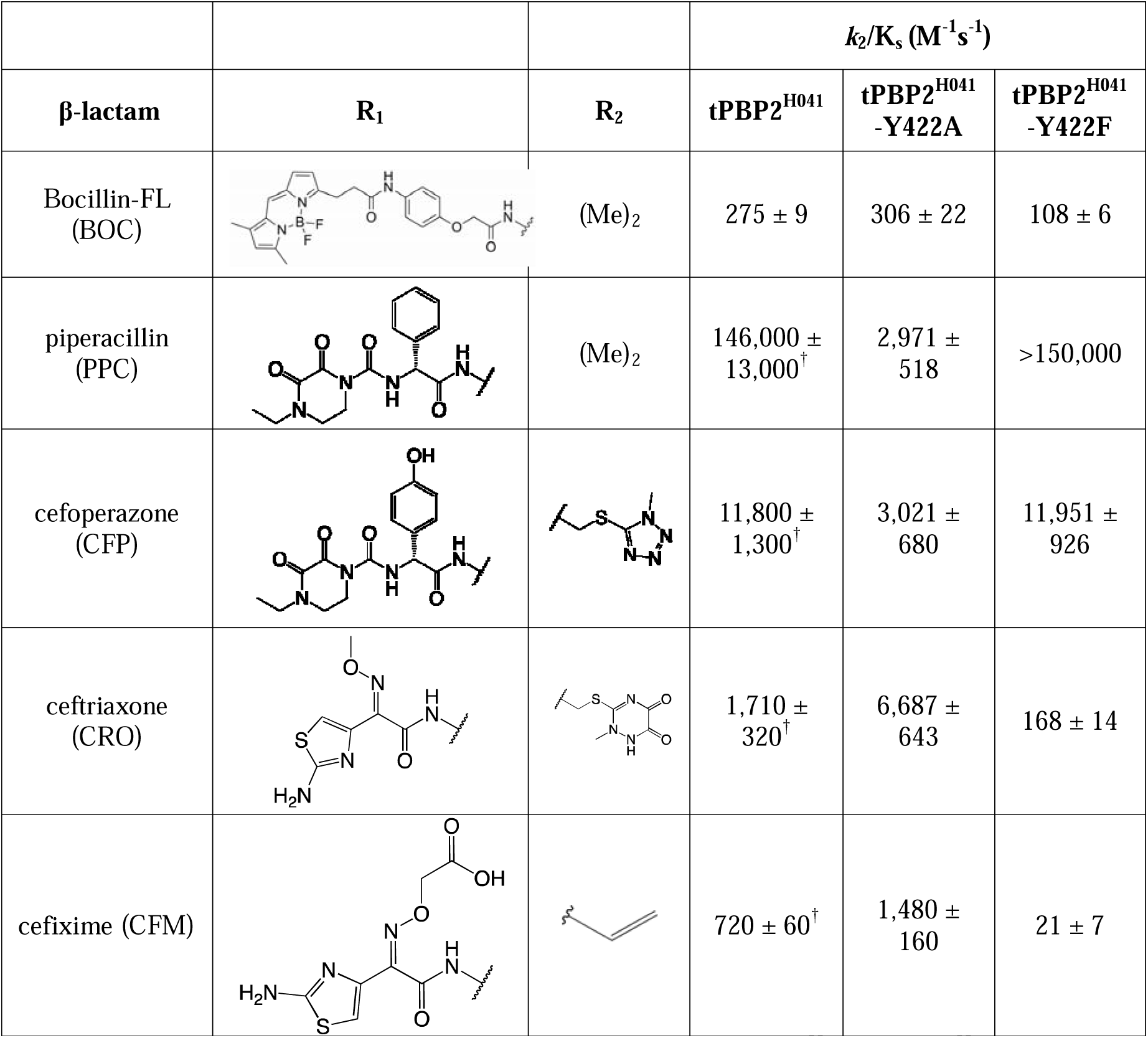
Acylation rates for selected β-lactams against tPBP2^H041^, tPBP2^H041^-Y422A and tPBP2^H041^-Y422F. ^†^Data previously reported by Turner *et al.* ^18^.

| β-lactam | $R_1$ | $R_2$ | $k_2/K_s$ ($M^{-1}s^{-1}$) | | |
| --- | --- | --- | --- | --- | --- |
|  |  |  | tPBP2<sup>H041</sup> | tPBP2<sup>H041</sup>-Y422A | tPBP2<sup>H041</sup>-Y422F |
| Bocillin-FL (BOC) | | (Me)<sub>2</sub> | 275 ± 9 | 306 ± 22 | 108 ± 6 |
| piperacillin (PPC) | | (Me)<sub>2</sub> | 146,000 ± 13,000<sup>†</sup> | 2,971 ± 518 | >150,000 |
| cefoperazone (CFP) | | | 11,800 ± 1,300<sup>†</sup> | 3,021 ± 680 | 11,951 ± 926 |
| ceftriaxone (CRO) | | | 1,710 ± 320<sup>†</sup> | 6,687 ± 643 | 168 ± 14 |
| cefixime (CFM) | | | 720 ± 60<sup>†</sup> | 1,480 ± 160 | 21 ± 7 |

### Thermostability of the acyl-enzyme complex correlates with acylation rate

We next investigated whether changes in thermostability of the acyl-enzyme complexes correlated with the acylation rate of the antibiotics for the Y422A and Y422F mutants of tPBP2^H041^. In the absence of a β-lactam, the melting temperature (T_m_) of the tPBP2^H041^-Y422A mutant was 5.3°C lower compared to tPBP2^H041^ (44.3 ± 0.3 vs 39.0 ± 0.0 °C) (Table 2), indicating that the tyrosyl ring inherently increases the thermostability of the protein. Since the T_m_ for tPBP2^H041^-Y422F was unchanged (44.6 ± 0.4°C), thermostability arises from the tyrosyl ring more than the hydroxyl.

**Table 2:**
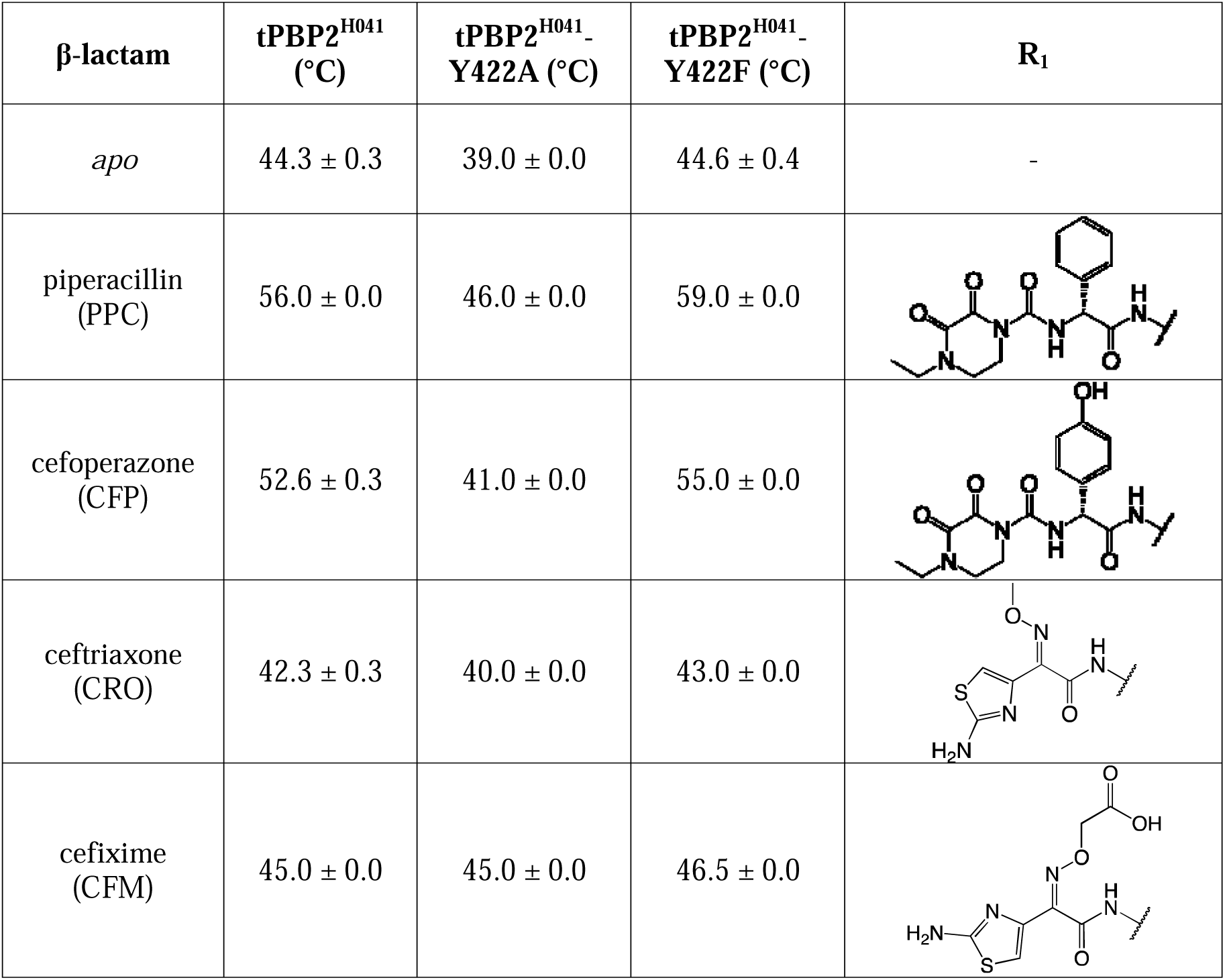
Thermal stabilities (T_m_) of tPBP2^H041^-Y422A acylated by a panel of β-lactams containing various R_1_ groups compared to acylated tPBP2^H041^.

| β-lactam | tPBP2<sup>H041</sup><br>(°C) | tPBP2<sup>H041</sup>-<br>Y422A (°C) | tPBP2<sup>H041</sup>-<br>Y422F (°C) | R<sub>1</sub> |
| --- | --- | --- | --- | --- |
| *apo* | 44.3 ± 0.3 | 39.0 ± 0.0 | 44.6 ± 0.4 | - |
| piperacillin<br>(PPC) | 56.0 ± 0.0 | 46.0 ± 0.0 | 59.0 ± 0.0 |  |
| cefoperazone<br>(CFP) | 52.6 ± 0.3 | 41.0 ± 0.0 | 55.0 ± 0.0 |  |
| ceftriaxone<br>(CRO) | 42.3 ± 0.3 | 40.0 ± 0.0 | 43.0 ± 0.0 |  |
| cefixime<br>(CFM) | 45.0 ± 0.0 | 45.0 ± 0.0 | 46.5 ± 0.0 |  |

We have previously observed that acylation by ureido β-lactams increases the T_m_ of tPBP2^H041^, while acylation by ceftriaxone slightly decreases it ^19^. Acylation by both piperacillin and cefoperazone similarly increased the thermostability of tPBP2^H041^-Y422A, but the magnitude of this stabilization was lower compared to tPBP2^H041^. Piperacillin increased the T_m_ of tPBP2^H041^ by 11.7°C ^19^, but only by 7.0°C for tPBP2^H041^-Y422A. The effect was more prominent with cefoperazone, going from an 8°C increase in T_m_ for tPBP2^H041^ to only a 2°C increase for the mutant. Ceftriaxone and cefixime both had little effect on T_m_ of tPBP2^H041^-Y422A, in comparison to a slight increase (2°C) when ceftriaxone was added to tPBP2^H041^. Thermostabilities for the Y422F mutant in the presence of the same antibiotics were similar to the wild-type protein, albeit with slightly higher T_m_ values. Taken together, these results suggest that the interaction of Tyr422 with the 2,3-dioxopiperazine ring of cefoperazone and piperacillin is stabilizing, consistent with their higher activities, whereas its interaction with the aminothiazole groups of ceftriaxone and cefixime contributes less.

### Mutation at position 422 does not alter the structure of tPBP2^H041^

Since Tyr422 constitutes a large, hydrophobic residue within the active site and its mutation to alanine both alters acylation activity and lowers the thermal stability of the protein, we next determined whether the structure of tPBP2^H041^-Y422A is altered compared to the unmutated protein. The structure was determined at 2.6 Å resolution with crystals obtained in the same P2_1_2_1_2_1_ system as tPBP2^H041 2^ (Table 3). It shows that mutation of Tyr422 to alanine does not alter the architecture of tPBP2^H041^, including the active site region and β3-β4 loop (Fig. S2). The two structures can be superimposed with an RMSD value of 0.22 Å for common Cα atoms. In addition, aside from the absence of a phenol ring side chain, there are no visible changes in structure around the mutation site. The same is true for the Y422F mutant, whose crystal structure was also solved at 2.6 Å resolution (Table 3). For both mutants, the absence of structural alterations beyond the mutation site in each case shows that kinetic differences can be attributed to the change in amino acid side chain.

**Table 3:** Crystallographic data collection and model refinement statistics for the tPBP2^H041^-Y422A *apo* and ceftriaxone (CRO)-acylated crystal structures, and tPBP2^H041^-Y422F *apo* and cefoperazone (CFP)-acylated crystal structures.

|  | **Y422A-*apo*** | **Y422F-*apo*** | **Y422A-CRO** | **Y422F-CFP** |
| --- | --- | --- | --- | --- |
| **Data collection** |  |  |  |  |
| Resolution range | 45.86–2.60<br>(2.64–2.60) | 39.23-2.60<br>(2.69-2.60) | 40.98–2.40<br>(2.44– 2.40) | 39.1-2.60<br>(2.64-2.60) |
| Space group | P2<sub>1</sub>2<sub>1</sub>2<sub>1</sub> | P2<sub>1</sub>2<sub>1</sub>2<sub>1</sub> | P2<sub>1</sub>2<sub>1</sub>2<sub>1</sub> | P2<sub>1</sub>2<sub>1</sub>2<sub>1</sub> |
| Unit cell *a*, *b*, *c* (Å) | 50.5, 61.2,<br>108.8 | 50.6, 61.5, 109.1 | 49.8, 59.5,<br>112.9 | 50.6, 61.6, 110.1 |
| Unique reflections | 10,931 (546) | 10,939 (1,055) | 13,460 (613) | 10,328 (548) |
| Multiplicity | 6.5 (6.2) | 11.6 (9.1) | 6.9 (5.7) | 6.8 (6.8) |
| Completeness (%) | 99.5 (99.5) | 99.9 (99.5) | 97.6 (90.3) | 92.6 (99.6) |
| Mean I/sigma(I) | 9.0 (2.1) | 12.6 (3.8) | 8.5 (2.0) | 16.5 (1.7) |
| Rmerge (%) | 16.6 (44.0) | 15.4 (38.1) | 13.0 (34.1) | 18.1 (47.0) |
| Rpim (%) | 6.9 (19.0) | 4.8 (13.2) | 5.2 (13.9) | 7.2 (18.5) |
| CC<sub>1/2</sub> | 0.975 (0.846) | 0.971 (0.899) | 0.985 (0.902) | 0.968 (0.909) |
| **Refinement** |  |  |  |  |
| Resolution range | 45.86–2.60<br>(2.66–2.60) | 39.23-2.46<br>2.52-2.46 | 40.98–2.40<br>(2.46–2.40) | 39.1-2.60<br>(2.67-2.60) |
| R-work (%) | 18.6 (25.4) | 20.8 (20.4) | 19.0 (26.5) | 22.1 (25.6) |
| R-free (%) | 25.3 (33.8) | 25.1 (38.6) | 24.3 (30.4) | 29.2 (35.6) |
| No. of non-hydrogen<br>protein atoms | 2,422 | 2,452 | 2,446 | 2,452 |
| No. of ligand atoms | - | - | 42 | 37 |
| No. of waters | 14 | 28 | 12 | 2 |
| RMSDs from ideal<br>stereochemistry: |  |  |  |  |
| RMS (bonds) | 0.007 | 0.006 | 0.007 | 0.007 |
| RMS (angles) | 1.70 | 1.71 | 1.93 | 1.70 |
| Ramachandran plot: |  |  |  |  |
| Ramachandran<br>favored (%) | 97.8 | 97.8 | 98.4 | 97.5 |
| Ramachandran<br>allowed (%) | 1.6 | 2.2 | 1.6 | 2.2 |
| Ramachandran<br>outliers (%) | 0.0 | 0.0 | 0.0 | 0.3 |
| B-factors: |  |  |  |  |
| Mean B-factor<br>(protein atoms) (Å<sup>2</sup>) | 27.3 | 24.1 | 35.4 | 42.8 |
| Ligand (Å<sup>2</sup>) | - | - | 44.3 | 49.4 |
| Waters (Å<sup>2</sup>) | 20.7 | 15.6 | 29.1 | 23.4 |
| PDB code | 9OSK | XX | 9OSL | XX |

### Crystal structure of ceftriaxone in complex with tPBP2^H041^-Y422A resembles ceftriaxone-acylated tPBP2^FA19^

To provide a framework to interpret the activity data, we determined the crystal structures of tPBP2^H041^-Y422A acylated by ceftriaxone (tPBP2^H041^-Y422A-CRO) at 2.4 Å resolution and of tPBP2^H041^-Y422A acylated by cefoperazone (tPBP2^H041^-Y422F-CFP) (Fig. 4 and Table 3) at 2.6 Å resolution. For both structures, a covalent bond is clearly visible between Ser310 and the β-lactam, confirming an acylated species (Fig. 4A & B). The majority of each cephalosporin molecule is visible in the |Fo|-Fc| electron density maps, except for weak density at the C3 vinyl of the dihydrothiazine ring and the carboxy methyloxime in the R_1_ group of ceftriaxone and the C3 vinyl and carboxylate of cefoperazone. The dioxo-triazine (ceftriaxone) and methyltetrazole (cefoperazone) leaving groups appear to have been eliminated from the C3 position following acylation, as expected ^35^.

**Fig. 4:**
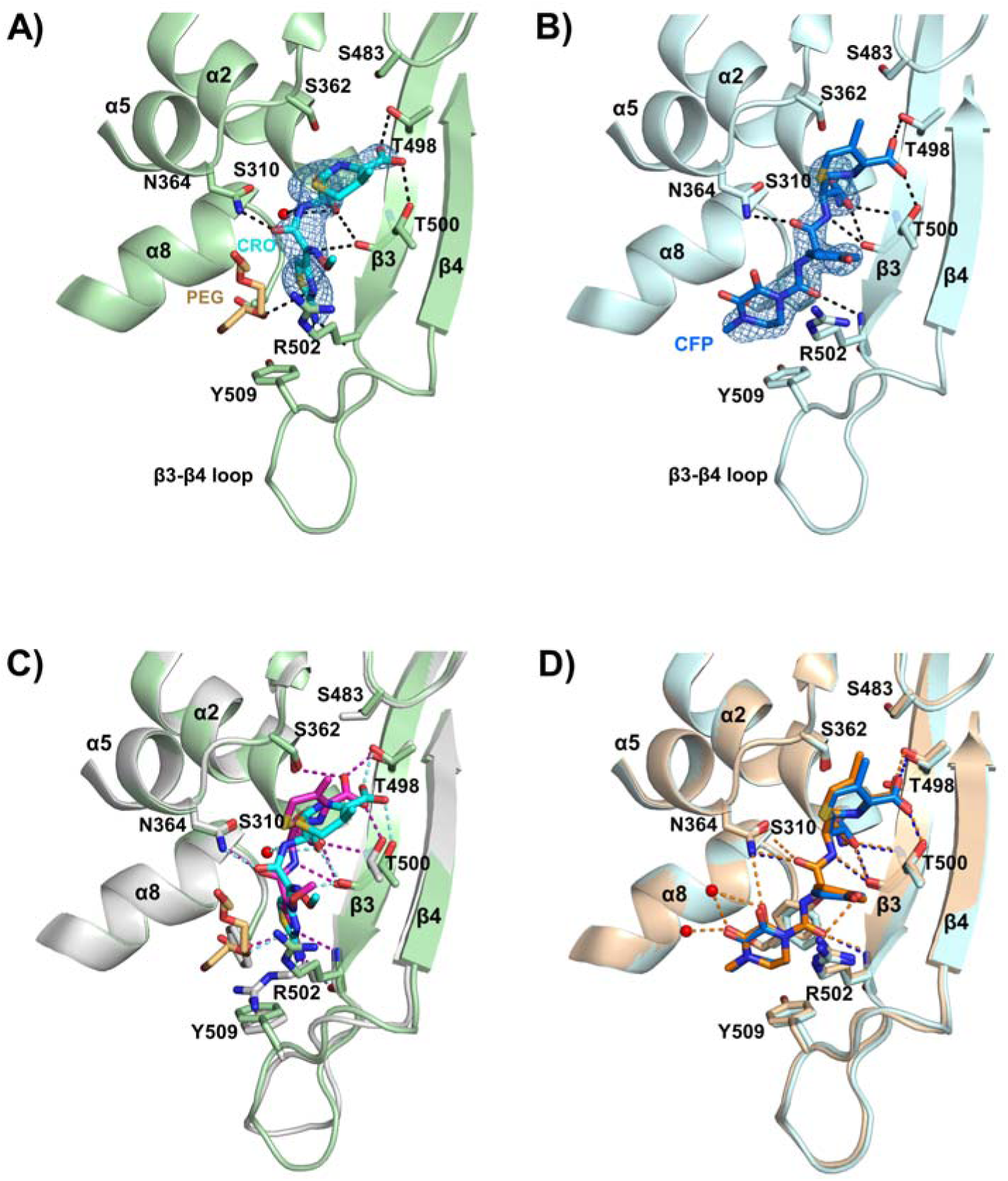
Structures of tPBP2^H041^ mutated at position 422. **A)** Structure of tPBP2^H041^-Y422A (green backbone) acylated by ceftriaxone (magenta bonds). **B)** Structure of tPBP2^H041^-Y422F acylated by cefoperazone (light blue backbone/blue bonds). **C)** Superimposition of tPBP2^H041^-Y422A and tPBP2^FA19^ structures (grey backbone/magenta bonds), each acylated by ceftriaxone. **D)** Superimposition of tPBP2^H041^-Y422F, and tPBP2^H041^ (wheat backbone/orange bonds) structures, each acylated by cefoperazone. For all panels, only the active site region is shown and potential hydrogen bonds are shown by dashed lines. |Fo|-|Fc| electron densities are shown as a blue mesh. Red spheres indicate ordered water molecules and a molecule of polyethylene glycol 600 (PEG) from the crystallization solution in the tPBP2^H041^-Y422A structure is colored with yellow bonds.

The *apo* and ceftriaxone-acylated structures of tPBP2^H041^-Y422A can be superimposed with RMSD value of 0.69 Å for common Cα atoms, showing a high similarity of the structures. In the active site, the R_1_ aminothiazole of ceftriaxone occupies the position previously held by Tyr422 in both *apo* and acylated structures of tPBP2^H041^. An interesting new feature is the presence of a molecule of polyethylene glycol (PEG) 600 from the crystallization solution that has bound in the active site adjacent to the aminothiazole group (Fig. 4A). Because a similar PEG is not observed in the ceftriaxone-acylated structure of tPBP2^H041 2^, its binding is likely due to the absence of the tyrosyl side chain at position 422.

A surprising outcome revealed by the crystal structure of tPBP2^H041^-Y422A-CRO is that the β3-β4 loop now adopts the same inward conformation seen in the structure of tPBP2^FA19^-CRO (Fig. 3B & C), in contrast to its outbent conformation in tPBP2^H041^-CRO. This is the first time an inward conformation for the loop has been observed with acylated structures of ceftriaxone in the background of tPBP2^H041^. Interestingly, the absence of Tyr422 means that this conformation is observed without the edge-π stack involving Tyr422, Phe504 and Tyr509 in tPBP2^FA19^-CRO, which was previously hypothesized to contribute to the stability of the inward conformation of the β3-β4 loop (Fig. S3) ^19^. An additional factor in tPBP2^H041^ may be that Phe504 is replaced by leucine, *i.e.* one of the two loop resistance mutations.

Because the β3-β4 loop occupies the inward position, polar contacts formed by ceftriaxone in tPBP2^H041^-Y422A-CRO are the same as those in tPBP2^FA19^-CRO (Fig. 4C). The side chain of Thr498 of the β3 strand is rotated to contact the β-lactam carboxylate, and the β3 strand itself is rotated toward Ser310 to form the canonical oxyanion hole in the same manner as in tPBP2^FA19 1^. In addition, there are similar interactions around the R_1_ group, as seen in tPBP2^FA19^-CRO ^1^. Overall, it can be concluded that tPBP2^H041^ adopts a tPBP2^FA19^-like binding mode for ceftriaxone when Tyr422 is mutated to an alanine.

As would be expected, the *apo* and cefoperazone structures of tPBP2^H041^-Y422F are very similar to one another, with an RMSD value of 0.46 Å for common Cα atoms. Commensurate with the high activity of piperacillin against this mutant (Table 1), the β3-β4 loop occupies the inward position and overall, the structure superimposes closely with the equivalent structure of tPBP2^H041^ (Fig. 4D). The absence of the tyrosyl side chain means that a hydrogen bond no longer forms with the amide group of cefoperazone but aside from this, the contacts are the same as seen in tPBP2^H041^ in complex with the same antibiotic.

### Mutation of Tyr422 is not viable in *N. gonorrhoeae*

Since Tyr422 impacts the acylation rates of β-lactams, we hypothesized that mutations at 422 may also affect the survival and fitness of *N. gonorrhoeae* due to disruption of the essential transpeptidation reaction. To examine this, we incorporated mutations into the *penA* allele of strain H041 (*penA41-Y422X*) and transformed the mutant constructs into the susceptible gonococcal strain, FA19. *N. gonorrhoeae* is naturally competent and will replace its endogenous *penA* allele with the mutant *penA41-Y422X* allele, provided that the encoded protein is capable of catalyzing essential transpeptidase activity at a level sufficient to support growth. Sub-MIC concentrations of cefixime were used to select for cells that had incorporated the *penA41* allele. Colonies that grew in the selection conditions were sequenced to detect the presence of mutations at position 422.

All of the transformants that grew on sub-MIC levels of cefixime had incorporated at least part of the *penA41* allele; however, none of these transformants had a mutation at position 422 (Fig. S4). In fact, some of the isolates had incorporated all of the *penA41* sequence but with Tyr422 intact. One possibility for why Y422X *penA41* mutants could not be isolated is that PBP2^H041^ is already moderately compromised in essential TPase activity ^36^, and incorporation of a Tyr422 mutation is sufficient to make the protein non-functional. Therefore, we also made Y422X mutations in the wild-type *penA* allele from FA19 (*penA19*) and incorporated a silent restriction site nearby to aid in identifying colonies that had recombined the mutant allele. Out of 72 colonies screened, only one incorporated the silent restriction site, but it still had a tyrosine at position 422. We then added a mutation (G545S) that modestly increases cefixime resistance to allow selection but minimize the effects on function. This increased our ability to identify transformants, but most isolates incorporated the G545S mutation without also incorporating the Y422X mutation and its associated silent restriction site. All of the colonies (6 in total) that did have the silent restriction site had a tyrosine codon at position 422. These results suggest that Tyr422 is essential for the TPase reaction catalyzed by PBP2, with mutations at this position disrupting viability.

To understand how Tyr422 might impact TPase activity, we modelled the pentapeptide portion of the Gram-negative peptidoglycan substrate into the electron density for cefoperazone and performed one round of crystallographic refinement against the dataset for the previously solved tPBP2^H041^-cefoperazone complex ^19^. The peptide fit surprisingly well into the cefoperazone density, with most atoms occupying density. The model indicates that the peptide is positioned antiparallel to β3 to become an extension of the β3-β11 sheet (Fig. 5). The branch point of the peptide matches the branch at the R_1_ group of cefoperazone (Fig. 5B) and the 2,3-diketopiperazine ring overlaps with the *iso*-Glu region of the peptide positioned above the sidechain of Tyr422 (as viewed in Fig. 5B). The central role that Tyr422 apparently plays in the positioning of the pentapeptide is consistent with mutations at this position being non-viable in *N. gonorrhoeae*.

**Fig. 5:**
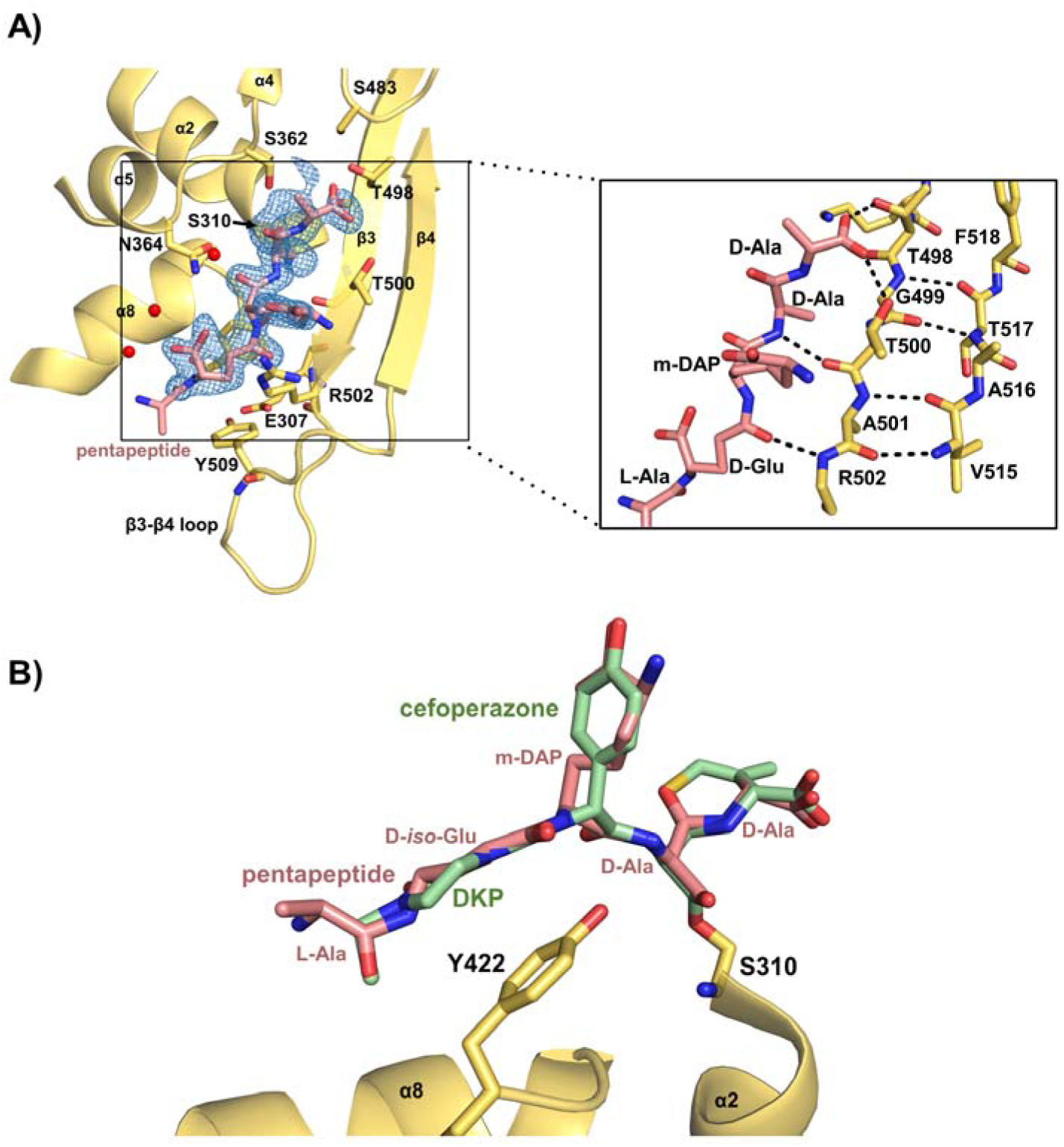
Peptidoglycan pentapeptide modeled into cefoperazone electron density. **A)** Peptidoglycan pentapeptide modeled into the |Fo|-|Fc| density (blue mesh) corresponding to cefoperazone. The inset shows how the modeled peptide is positioned alongside the β3 and β4 strands, thereby extending the β sheet of the transpeptidase domain. **B)** Superimposition of tPBP2^H041^ acylated by cefoperazone and modeled pentapeptide showing the proximity of Tyr422 to the D-*iso*Glu and diketopiperazine (DKP) groups. In all views, tPBP2^H041^ is represented by a yellow cartoon or bonds. The peptide is colored pink and cefoperazone is green.

### Tyr422 is conserved in over half of structurally related PBPs

With transformation experiments and modelling suggesting an essential role for Tyr422 in transpeptidation, we examined the degree of conservation of this residue across other Class B PBPs that share homology with PBP2. Using PDB code 6P53 as the search term, we accessed the homology-derived secondary structure of proteins (HSSP) multiple sequence alignment database ^31^ and calculated the percent identity for residues at the equivalent position to 422. A tyrosine was observed at this position in 55% of the 224 aligned sequences, with alternative residues being predominantly histamine and glutamine (Fig. 6). Interestingly, high conservation for two flanking glycines is observed (residues 421 and 423 using NgPBP2 numbering). These glycines may impart flexibility to position 422 that is necessary for transpeptidase function.

**Fig. 6:**
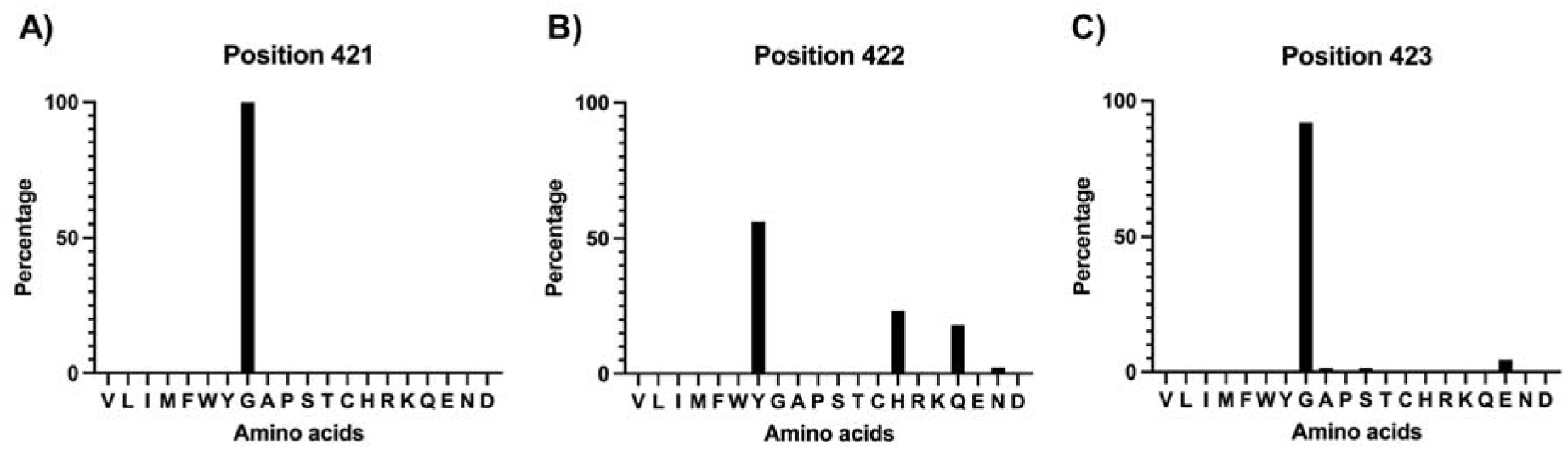
Conservation of tyrosine 422 and adjacent residues. The MRS server was used to fetch aligned sequences of Class B PBPs homologous to PBP2 from *N. gonorrhoeae* from the Homology Secondary Structure of Proteins (HSSP) databank using the PDB code 6P53 (tPBP2^FA19^) as the search term. Of the 1,013 sequences aligned, 224 sequences with unique genera were selected for calculating percent amino acid conservation at each position 422 and adjacent residues. Graphs were prepared using GraphPad Prism 8.0.2.

To examine the conservation of Tyr422 within *N. gonorrhoeae*, 800 unique *penA* loci isolates in the PubMLST database ^33^ were aligned. 100% of loci with intact *penA* coding regions had tyrosine at the equivalent of position Tyr422. When this alignment was expanded to include all *Neisseria* species available in the database (including *N. meningitidis* and 40 other *Neisseria* species), all 5,274 loci had an equivalent Tyr422 amino acid.

### Equivalent interactions between Tyr409 and β-lactam R_1_ groups occur in crystal structures of *Pseudomonas aeruginosa* PBP3

*Pseudomonas aeruginosa* PBP3 (*Pa*PBP3) shares 38% sequence identity with *N. gonorrhoeae* PBP2, with a tyrosine residue (Tyr409) found at the equivalent position to Tyr422. Structures of acylated *Pa*PBP3 show similar interactions of the β-lactam R_1_ group with Tyr409 (Fig. 7) as those in tPBP2^FA19^ with Tyr422. In the structure of ceftazidime-acylated PBP3 ^37^ (PDB: 3PBO), Tyr409 is displaced from its position in the *apo* structure in order to accommodate the aminothiazole R_1_ group, similar to Tyr422 in the ceftriaxone-acylated structure of tPBP2^FA19^ (Fig. 7A, C). However, the magnitude of this displacement is greater in *Pa*PBP3 than in equivalent tPBP2^FA19^ structures, with an approximately 104° rotation of the side chain that results in a 9 Å shift of the tyrosine hydroxyl from its starting position. The shift of the tyrosyl side chain from *apo* to ceftriaxone-acylated tPBP2^FA19^ is relatively minor in comparison, rotating ∼45° to accommodate the aminothiazole group (see Fig. 2B). Despite the different positions of the tyrosine side chain, the two aminothiazole groups are observed to occupy the same position.

**Fig. 7:**
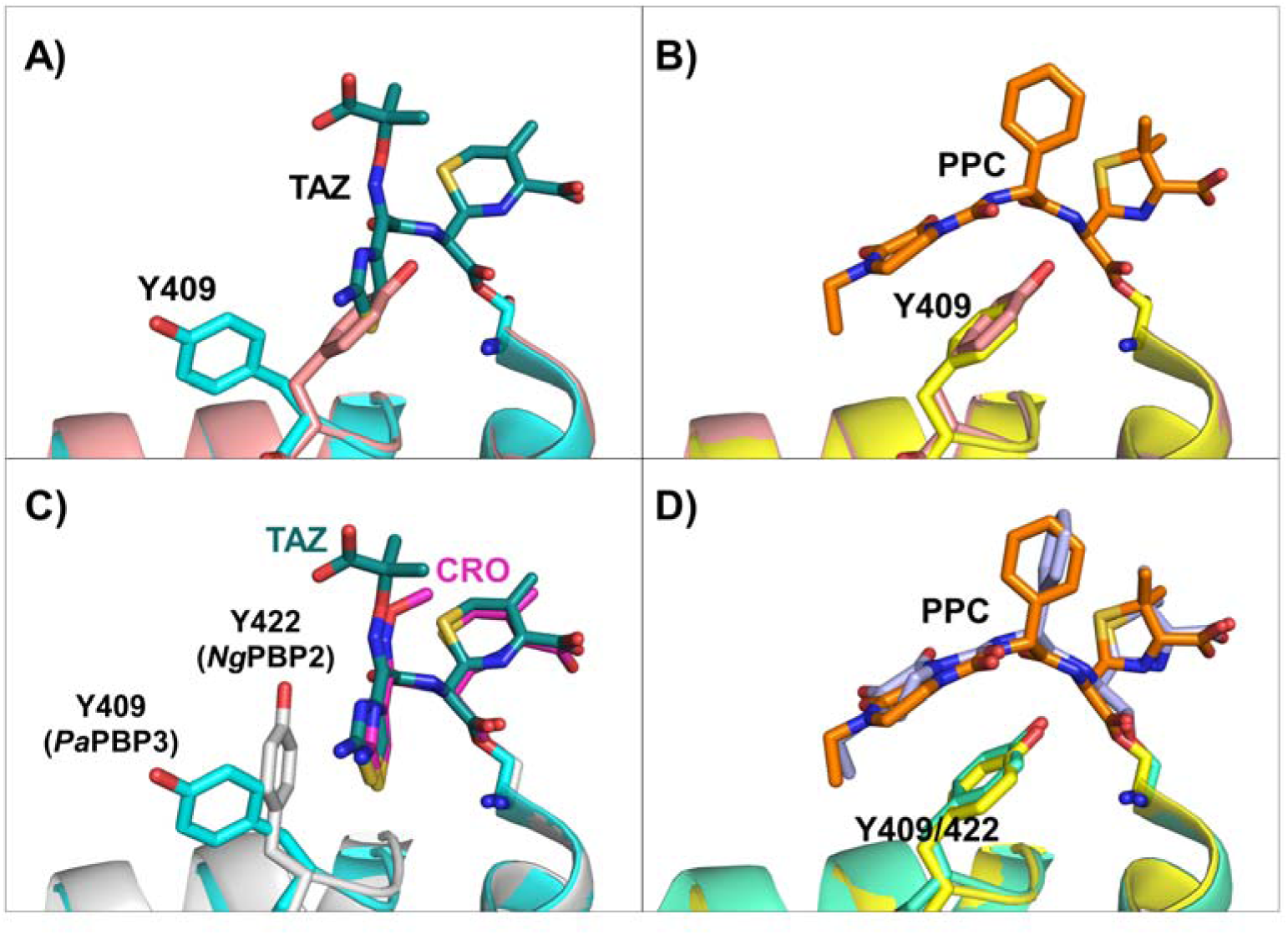
Comparison of Pseudomonas aeruginosa PBP3 (PaPBP3) with N. gonorrhoeae PBP2. **A)** Ceftazidime (TAZ)-acylated *Pa*PBP3 (cyan backbone/dark teal bonds) superimposed onto *apo Pa*PBP3 (salmon backbone). **B)** Piperacillin (PPC)-acylated *Pa*PBP3 (yellow backbone/orange bonds) superimposed onto *apo Pa*PBP3 (salmon backbone). **C)** Ceftazidime-acylated *Pa*PBP3 (cyan backbone/dark teal bonds) superimposed onto ceftriaxone (CRO)-acylated tPBP2^FA19^ (gray backbone/magenta bonds). **D)** Piperacillin-acylated *Pa*PBP3 (yellow backbone/orange bonds) superimposed onto piperacillin-acylated tPBP2^FA19^ (green backbone/light blue bonds).

In contrast to the situation with ceftriaxone, Tyr409 does not change position when going from an *apo* form to being acylated by piperacillin and the 2,3-diketopiperazine ring stacks over Tyr409 in the active site (Fig. 7B). This arrangement is essentially the same as that seen in the piperacillin-acylated structure of tPBP2^FA19^ (Fig. 7D) and suggests that the difference in binding of β-lactams containing an aminothiazole versus diketopiperazine R_1_ group might occur more generally in PBPs. Additionally, the equivalent of the β3-β4 loop in *Pa*PBP3 is disordered in the ceftazidime-acylated structure but fully ordered in an inward conformation when acylated by piperacillin. These observations suggest Tyr409 interacting with the aminothiazole group in *Pa*PBP3 can also affect the conformation of the loop and may also be important for acylation.

## Discussion

The continued emergence and spread of cephalosporin-resistant *N. gonorrhoeae* highlights the need to understand the molecular determinants governing inhibition of PBP2 by β-lactam antibiotics. Tyr422 is a highly conserved active-site residue positioned adjacent to the R_1_ substituent of β-lactams, yet its role in acylation was unclear. This study was prompted by the different positions of the R_1_ aminothiazole group of ceftriaxone with respect to the Tyr422 side chain in structures of tPBP2^FA19^ and tPBP2^H041 1, 2^. Our kinetic, structural, and genetic data demonstrate that Tyr422 contributes to both β-lactam inhibition and transpeptidase function, but in a manner that depends strongly on the nature of the ligand. The data also reveal that Tyr422, through functional coupling with the β3-β4 loop, influences formation of the high-activity state for acylation. Hence, there appears an unexpected mechanistic connection between an active-site recognition residue and a structural element of PBP2 that is associated with resistance.

A major finding of this study is that Tyr422 influences acylation as predicted, but with opposite effects depending on the identity of the β-lactam side chain. The Y422A substitution increased acylation rates for ceftriaxone and cefixime but substantially reduced acylation rates for cefoperazone and piperacillin, with the latter exhibiting an approximately 120-fold decrease. The apparent paradox of Tyr422 both promoting or inhibiting activity suggests that its contribution extends beyond simple packing interactions with the R_1_ group but involves structural organization of the active site. This is also supported by the kinetic effects being accompanied by corresponding changes in thermostability of the acyl-enzyme complexes, where higher thermostability correlates with higher activity, suggesting that the thermostability increase results from reorganization of the active site in addition to the interaction between Tyr422 and the R_1_ group.

The structure of tPBP2^H041^-Y422A acylated by ceftriaxone provides insight into the molecular basis of this behavior. In contrast to the tPBP2^H041^-ceftriaxone complex, the β3-β4 loop adopts an inward conformation in the mutant and Thr498 rotates to engage the ceftriaxone carboxylate. This arrangement closely resembles the conformation observed in tPBP2^FA19^ when acylated by β-lactams and is associated with higher acylation activity. In the tPBP2^H041^-ceftriaxone complex, the β3-β4 loop remains outbent and Thr498 is hydrogen bonded to Ser545 rather than the β-lactam carboxylate. The increase in ceftriaxone acylation observed upon removal of the Tyr422 side chain is therefore coupled with the conformational state of the β3-β4 loop associated with efficient acylation. Because the opposite effect is seen with cefoperazone and piperacillin, the findings cannot be explained solely by ligand binding. Instead, it appears that Tyr422 influences the β3-β4 loop through an allosteric coupling mechanism, thereby affecting formation of a productive acylation complex.

The influence of Tyr422 can be described in terms of the following model linking its interaction with the R_1_ group to the conformation of the β3-β4 loop and corresponding switching between high– or low-activity states for acylation. For β-lactams with relatively smaller R_1_ groups like ceftriaxone and cefixime, the interaction with Tyr422 is relatively weak, the β3-β4 loop remains outbent, and the protein remains in the lower-activity state. For piperacillin and cefoperazone, which have larger R1 groups, interaction with Tyr422 is stronger, thus favoring the inward conformation the β3-β4 loop and the higher-activity state. This means that Tyr422 is neither intrinsically beneficial or detrimental for acylation activity, but instead its effect is governed by whether the β-lactam ligand can harvest the energetic potential of the interaction to alter the conformation of the loop.

The behavior of ceftriaxone and cefixime vs. cefoperazone and piperacillin can now be rationalized within the context of the resistance-associated mutations present in H041. Previous studies established that substitutions in PBP2, including F504L and N512Y on the β3-β4 loop, contribute to reduced susceptibility by lowering rates of acylation. These are believed to alter the conformational accessibility of the high-activity state by the favoring the outbent conformation of the β3-β4 loop. In the presence of ceftriaxone or cefixime, whose aminothiazole R_1_ groups provide relatively limited interaction surfaces with Tyr422, the energetic contribution of ligand binding may be insufficient to favor the inward-loop conformation. Consequently, PBP2 is more likely to occupy the outbent state associated with lower acylation activity. In contrast, β-lactams containing larger R_1_ groups, such as the diketopiperazine groups of piperacillin and cefoperazone, interact more extensively with Tyr422, thus providing the energetic driving force to overcome the conformational restrictions imposed by the H041 resistance mutations and favoring the inward-loop state associated with efficient acylation. Consistent with this, mutation of Tyr422 to alanine markedly reduced acylation rates for piperacillin and cefoperazone and decreased thermostability of the corresponding acyl-enzyme complexes, whereas mutation to phenylalanine had minimal effect. Overall, the available kinetic and structural data support a model in which interactions of R_1_ groups with Tyr422 influence β3-β4 loop positioning and subsequent acylation efficiency, where the equilibrium between states is altered by resistance mutations.

The transformation experiments further indicate that Tyr422 plays an important role in the physiological function of PBP2. Substitutions at this highly conserved position were not tolerated in *N. gonorrhoeae*, suggesting that Tyr422 contributes directly to transpeptidase activity. Consistent with this interpretation, modeling of a peptidoglycan stem peptide into density corresponding to cefoperazone suggests that Tyr422 interacts with the *iso*-Glu residue of the natural substrate. If so, the interactions observed between Tyr422 and β-lactam R_1_ groups may represent exploitation of a recognition mechanism that evolved for substrate binding. β-lactams possessing extended R_1_ substituents achieve enhanced activity fortuitously because they more effectively mimic interactions formed with the natural substrate and alter the conformational equilibrium of the β3-β4 loop toward the inward state. This can also be viewed as Tyr422 acting as a “checkpoint” to monitor for ligands capable of stabilizing a productive active-site geometry.

The present findings have implications for the development of future PBP2 inhibitors. Third– and fourth-generation cephalosporins commonly contain aminothiazole R_1_ groups that enhance activity against Gram-negative pathogens, yet our results indicate that these substituents interact differently with susceptible and resistant forms of PBP2. In tPBP2^FA19^, the aminothiazole group is positioned inside Tyr422 and is associated with the productive inward-loop conformation. In tPBP2^H041^, however, the aminothiazole group is outside Tyr422 and appears less effective at promoting the same active-site arrangement. By contrast, larger R_1_ substituents, such as those present in cefoperazone and piperacillin, appear capable of engaging Tyr422 more effectively and may partially compensate for the conformational restrictions imposed by resistance mutations. This interpretation is consistent with our previous observation that cephalosporins containing larger aliphatic R_1_ groups exhibit enhanced acylation of tPBP2^H041^. Expanding the R_1_ substituent to exploit the region adjacent to Tyr422 therefore represents one potential strategy for improving activity against resistant gonococcal strains.

An alternative strategy may be to develop compounds that avoid dependence on interactions with Tyr422 altogether. Modifications at the C3 position can substantially influence acylation rates, as illustrated by the higher activity of ceftriaxone relative to cefixime. Similarly, carbapenems may be less affected by the conformational constraints associated with Tyr422 because they lack the extended R_1_ architecture. The successful use of ertapenem to treat ceftriaxone-resistant gonorrhea is consistent ^38^ with the possibility that carbapenems circumvent some of the mechanistic limitations imposed by the Tyr422/β3-β4 loop network.

In summary, our results demonstrate that Tyr422 is an important determinant of both β-lactam acylation and transpeptidase function. The effects of Tyr422 cannot be explained solely by direct interactions with antibiotic side chains but it appears that this residue additionally serves as a ligand-responsive conformational gate that links recognition of peptide substrate and the R_1_ groups of β-lactams with the position of the β3-β4 loop. In resistant PBP2 variants, this relationship becomes an important determinant of acylation efficiency and provides a mechanistic framework for understanding why β-lactams bearing different R_1_ substituents exhibit markedly different activities. These findings also identify a previously unrecognized aspect of PBP2 function and suggest new approaches for the design of β-lactams active against antimicrobial-resistant *N. gonorrhoeae*.

## Accession Codes

*N. gonorrhoeae* PBP2: Q8RR30 (UniProt)

Coordinates and structure factors have been deposited with the Protein Data Bank as follows:

- tPBP2^H041^-Y422A in *apo* form: 9OSK
- tPBP2^H041^-Y422A acylated by ceftriaxone: 9OSL
- tPBP2^H041^-Y422FA in *apo* form: *pending*
- tPBP2^H041^-Y422F acylated by cefoperazone: *pending*
- tPBP2^FA19^ acylated by piperacillin: 9OTG

## Supporting Information

**Supplemental Fig. 1:** Second-order rate of acylation of Bocillin-FL against the tPBP2^H041^-Y422A mutant

**Supplemental Fig. 2:** Superimposition of the crystal structures of tPBP2^H041^-Y422A and tPBP2^H041^

**Supplemental Fig. 3:** Superimposition of the crystal structures of tPBP2^FA19^ and tPBP2^H041^-Y422A acylated by ceftriaxone

**Supplemental Fig. 4:** Sequencing of *penA* of *N. gonorrhoeae* FA19 transformed with *penA^H041^* alleles mutated at position 422

**Supplemental Table 1:** PCR primers used for site-directed mutagenesis

**Supplemental Table 2:** Crystallographic data collection and model refinement statistics for the crystal structure of tPBP2^FA19^ acylated by piperacillin

## Acknowledgements

This work was supported by National Institutes of Health award AI164794 (to C.D. and R.A.N.) and U19 AI113170 and AI153521 (to R.A.N.). Use of the Advanced Photon Source was supported by the U.S. Department of Energy, Office of Science, Office of Basic Energy Sciences, under Contract No. W-31-109-ENG-38. Data were collected at Southeast Regional Collaborative Access Team (SER-CAT) 22-ID beam line at the Advanced Photon Source, Argonne National Laboratory. Supporting institutions may be found at www.sercat.org/members.html.

## Author contributions statement

CMS, RAN, and CD conceived the experiments; CMS, SB and MMB performed the experiments; CMS wrote the first draft of the manuscript, followed by editing and writing by CD, RAN, CMS, and MMB. All authors reviewed the manuscript. CD and RAN obtained funding for the work.

