## Supplemental Data for "Energetic coupling of an active site residue in penicillin-binding protein 2 from *Neisseria gonorrhoeae* with a resistance-associated conformational switch in the β3-β4 loop"

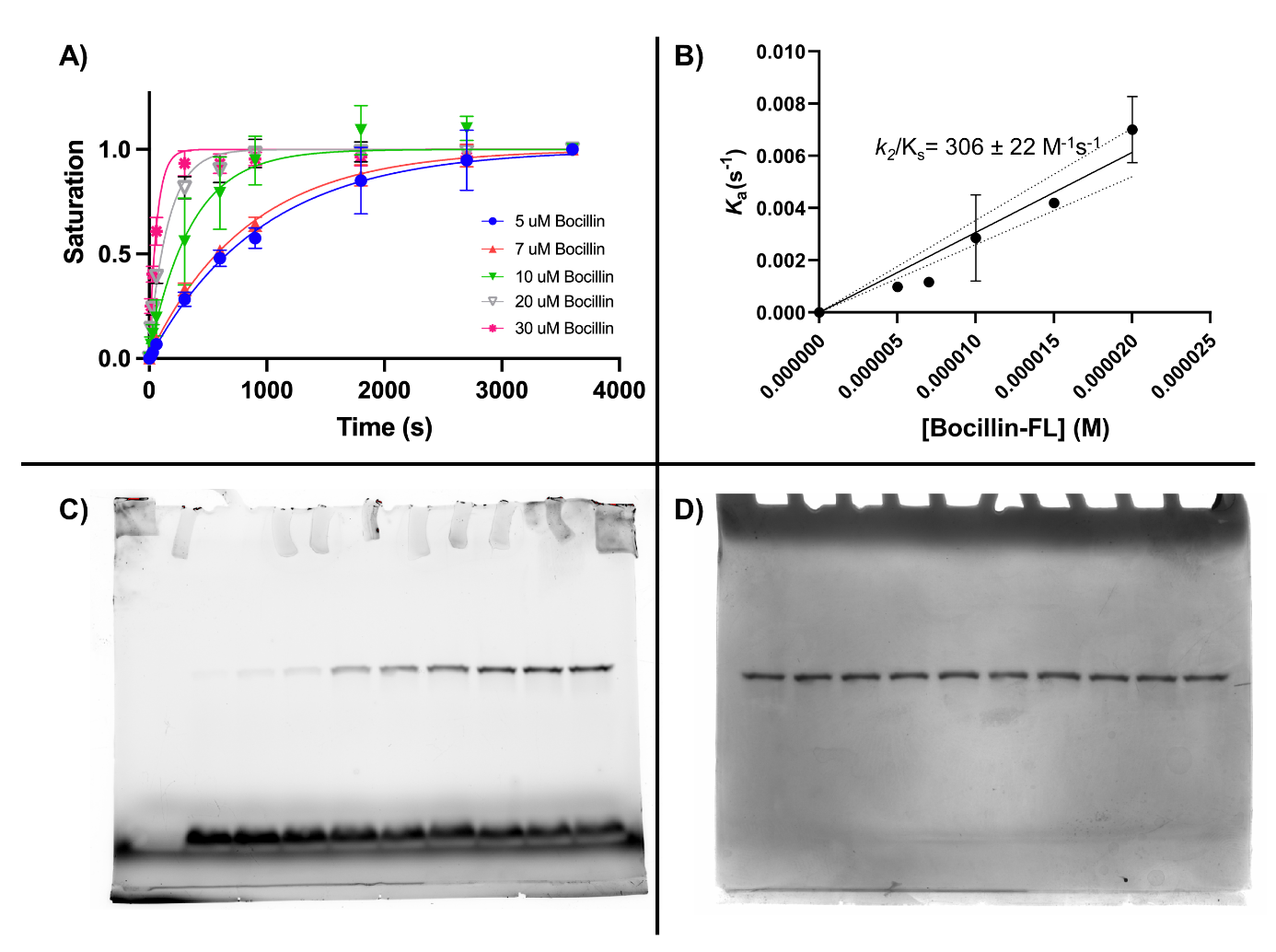

**Fig. S1: Second-order rate of acylation of Bocillin-FL against the tPBP2^H041^-Y422A** **mutant.** **A)** Pseudo-first order kinetics of tPBP2^H041^-Y422A at different Bocillin-FL concentrations. **B)** Plot of *k_a_* values against Bocillin-FL concentrations. The slope of the line is the calculated second-order acylation rate (*k*_2_/K_s_). **C)** Representative fluorescent image of an SDS-PAGE gel of tPBP2^H041^-Y422A acylated by Bocillin-FL over time. **D)** Coomassie stain for protein of a representative SDS-PAGE gel, showing equal amounts of protein in each condition.

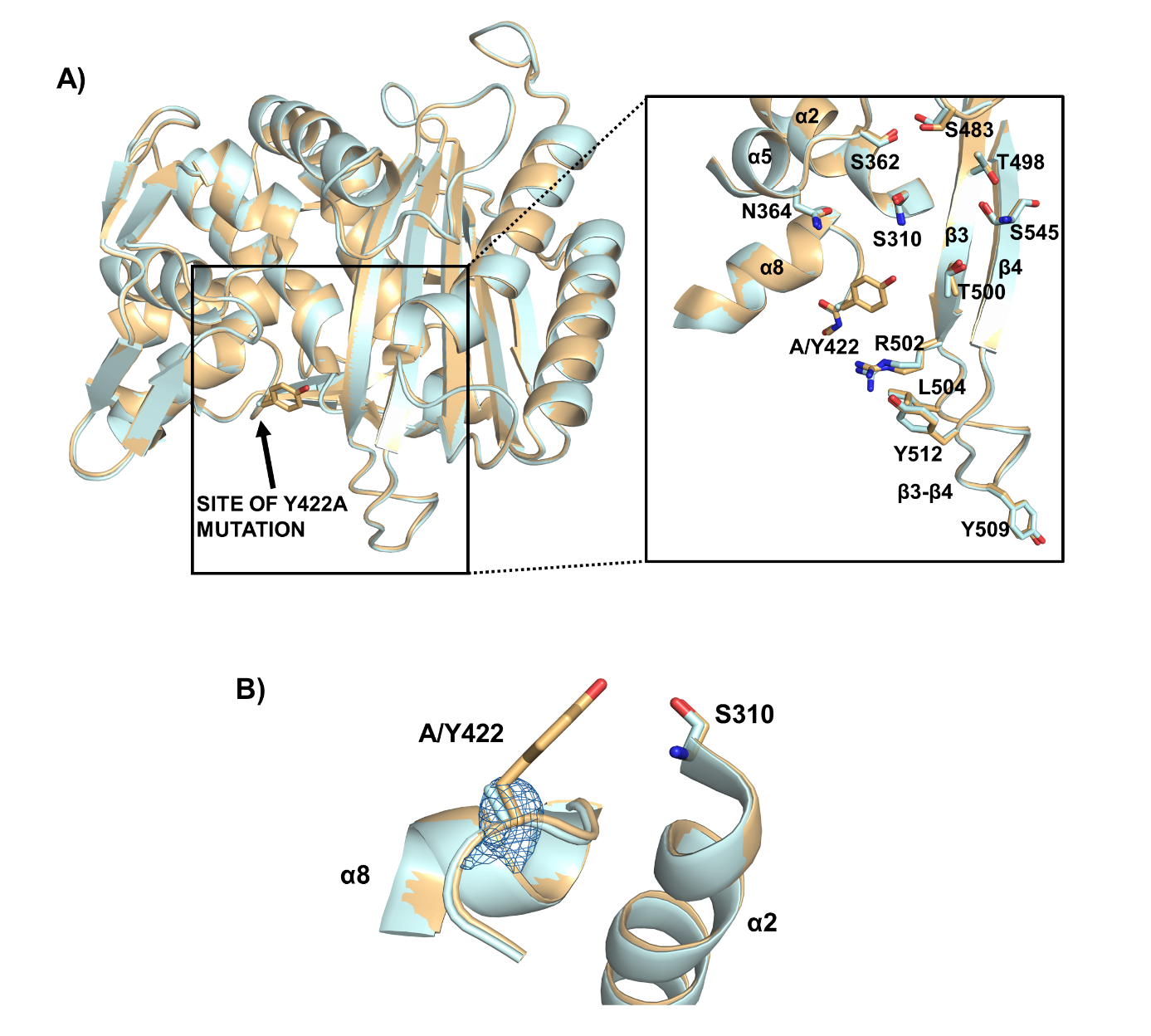

**Fig. S2: The Y422A mutation does not alter the structure of tPBP2^H041^**. **A)** Superimposition of the folds of tPBP2^H041^-Y422A in light blue and tPBP2^H041^ in light orange. Inset shows the active site region including the mutation site. **B)** The |Fo|-|Fc| density at position 422 in the tPBP2^H041^-Y422A structure, showing the absence of the tyrosyl side chain

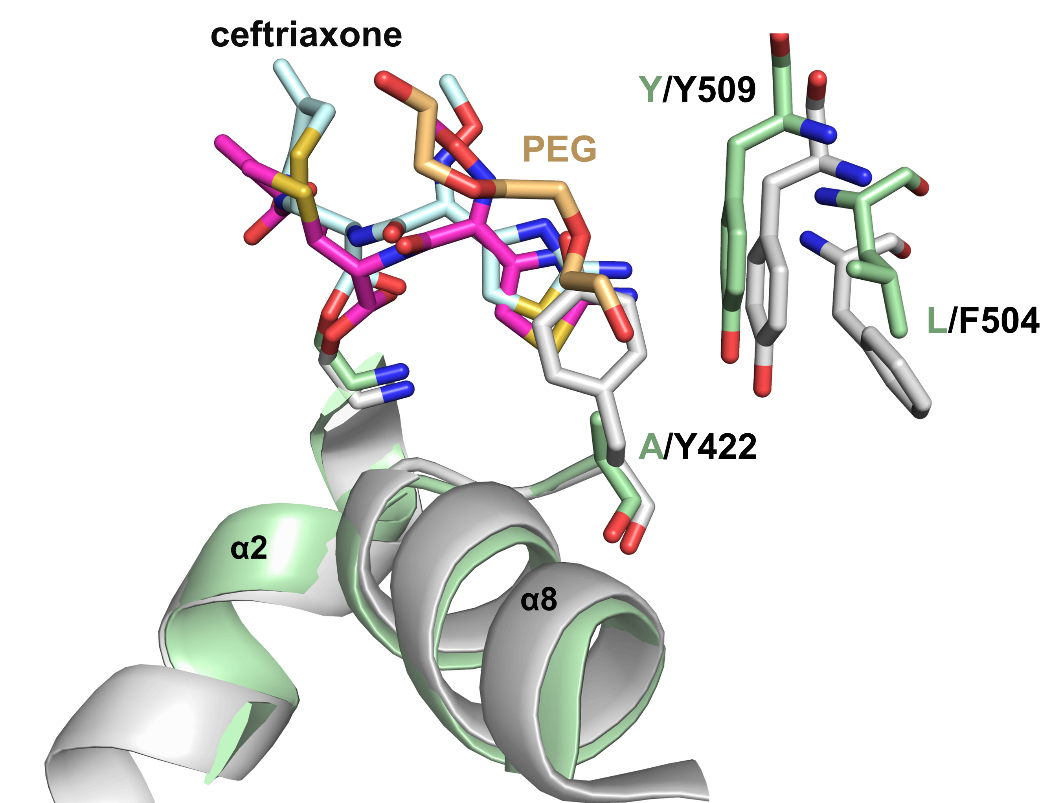

**Fig. S3: Inward conformation of the β3-β4 loop occurs in absence of edge-π stacking.** Structure of ceftriaxone-acylated tPBP2^FA19^ in white backbone and magenta bonds, in which edge-π stacking involving Tyr422, Phe504 and Tyr509 was observed. Structure of tPBP2^H041^-Y422A acylated by ceftriaxone, shown in light green backbone with cyan bonds, showing the absence of similar stacking due to the absence of the tyrosyl side chain. Note that in tPBP2^H041^, Phe504 is replaced by a leucine.

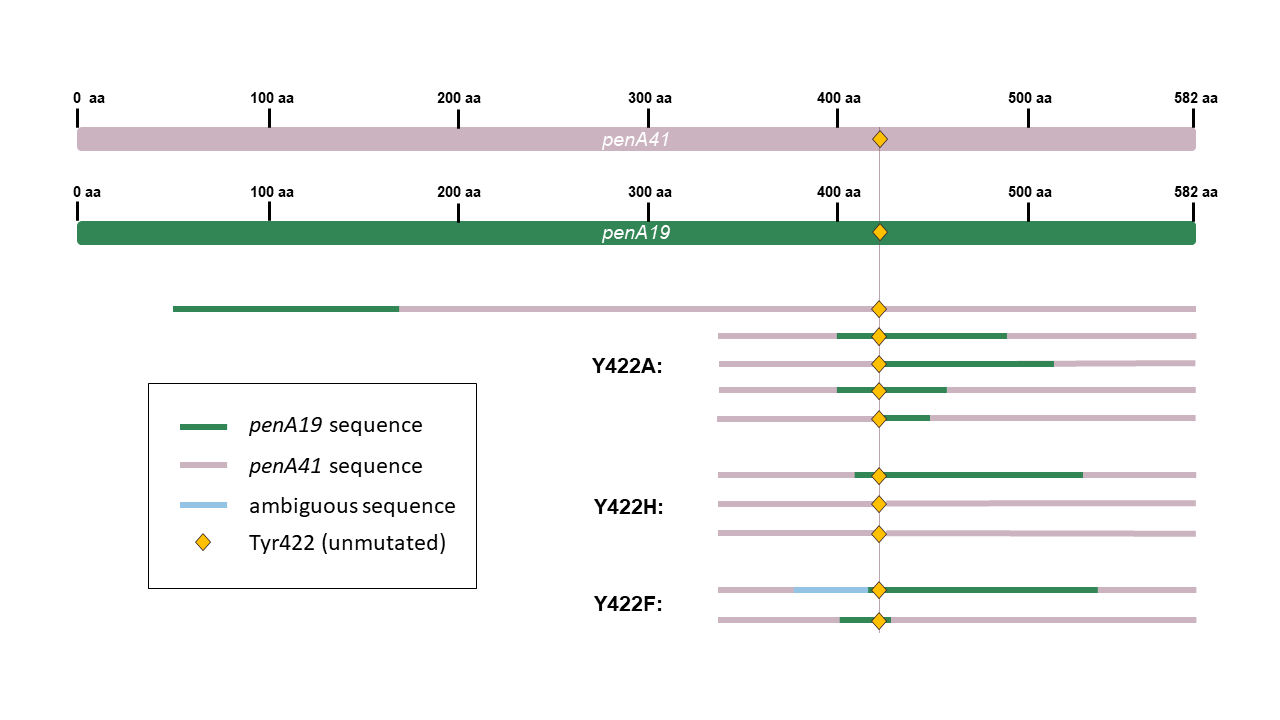

**Fig. S4: Y422X mutations cannot be introduced into FA19 via transformation.** Sequencing of individual transformants of FA19 transformed with *penA^H041^* alleles containing the indicated Tyr422 mutations. Full sequences of *penA41* (rose) and *penA^FA19^* (green) are shown at the top, with the individual sequences of the transformants shown below. The origins of the transformant sequences are color-coded to indicate from which allele (either penA41 or *penA^FA19^*) the sequence is derived. Ambiguous sequence (arising from a potentially a mixed colony) is shown in light blue. The location of Tyr422 (present in both *penA41* and *penA^FA19^*) is depicted by the orange diamond.

**Table S1.** *PCR* p*rimers used for site-directed mutagenesis*.

| Primer Name | Primer Sequence | Notes |
| --- | --- | --- |
| 5’ *penA*_45aa_Uptake | 5’- GCCGTCTGAACGCGGGCTGTATCTGCAGACGGTAA-3’ | Can be used for both *penA19* and *penA41* |
| 5’ *penA*_45aa | 5’- CGCGGGCTGTATCTGCAG-3’ | 5’ *penA*_45aa_Uptake without the *N. gonorrhoeae* uptake sequence. Can be used for both *penA19* and *penA41* |
| 3’ *murE*_31aa | 5’-CGGCTGTCTGAATGCAACAA-3’ |  |
| 5' *penA41*_Y422A | 5’-ACGATGTCTTTCGGTgccGGCCTGCAATTAAGC-3’ | Lowercase nucleotides indicate altered sequence |
| 3' *penA41_*Y422A | 5’-GCTTAATTGCAGGCCggcACCGAAAGACATCGT-3’ | Lowercase nucleotides indicate altered sequence |
| 5' *penA41_*Y422H | 5’-GACGATGTCTTTCGGTcATGGCCTGCAATTAAG-3’ | Lowercase nucleotide indicates altered sequence |
| 3' *penA41_*Y422H | 5’-CTTAATTGCAGGCCATgACCGAAAGACATCGTC-3’ | Lowercase nucleotide indicates altered sequence |
| 5' *penA41_*Y422F | 5’- GACGATGTCTTTCGGTTtTGGCCTGCAATTAAGCC-3’ | Lowercase nucleotide indicates altered sequence |
| 3' *penA41_*Y422F | 5’-GGCTTAATTGCAGGCCAaAACCGAAAGACATCGTC-3’ | Lowercase nucleotide indicates altered sequence |
| 5' *penA19_*Y422X_AccI | 5’-CAATTGAGCCTGCTGCAATTG-3’ | Used for both *penA19*-*Y422X+G545S* and *penA19*-*Y422H+G545S* |
| 3' *penA19_*Y422X_AccI | 5’-CAATTGCAGCAGGCTCAATTGtAGACCnnnACCGAAAGACATCGTCG-3’ | Lowercase nucleotides indicate altered sequence, “n” indicates random nucleotides |
| 3' *penA19_*Y422H_AccI | 5’-CAATTGCAGCAGGCTCAATTGtAGACCGTgACCGAAAGACATCGTCG-3’ | Lowercase nucleotides indicate altered sequence |
| 5’ *penA19*_G545S_Fse | 5’-ATTACaGCGGCGTAGTGGCcGGcCCGCCCTTCAAAAAAATTATGGGCGGC-3’ | Lowercase nucleotides indicate altered sequence |
| 3’ *penA19*_G545S_Fse | 5’- CGGgCCgGCCACTACGCCGCtGTAATAGCCGTTGGCAGTCGGTT-3’ | Lowercase nucleotides indicate altered sequence |
| 5’ *penA41*_317aa | 5’-ATTGCCAAAGCGCTGGATTC-3’ | Used for *penA41* mutant sequencing |
| 5' *penAwt_*317aa | 5’-ATTGCGAAGGCATTGGATG-3’ | Used for *penA19* mutant sequencing |

**Table S2.** *Crystallographic data collection and model refinement statistics for the crystal structure of tPBP2^FA19^ acylated by piperacillin.*

|  | tPBP2^FA19^-PPC |
| --- | --- |
| **Data collection** |  |
| Resolution range | 39.0 – 3.15 (3.26 – 3.15) |
| Space group | P 2_1_ |
| Unit cell *a, b, c* (Å), *α, β, γ* (°) | 45.0, 78.0, 87.2  90, 90.8, 90 |
| Unique reflections | 10,473 (1,053) |
| Multiplicity | 3.9 (4.0) |
| Completeness (%) | 99.0 (99.5) |
| Mean I/sigma(I) | 11.9 (3.3) |
| R-merge (%) | 12.8 (66.8) |
| R-pim (%) | 7.5 (39.2) |
| CC_1/2_ | 0.992 (0.731) |
| **Refinement** |  |
| Resolution range | 39.0 – 3.15 (3.23 – 3.15) |
| R-work (%) | 15.3 (23.3) |
| R-free (%) | 26.6 (28.8) |
| No. of non-hydrogen protein atoms | 4,952 |
| No. of ligand atoms | 72 |
| RMSDs from ideal stereochemistry: |  |
| RMS (bonds) | 0.004 |
| RMS (angles) | 1.38 |
| Ramachandran plot: |  |
| Ramachandran favored (%) | 91.5 |
| Ramachandran allowed (%) | 98.6 |
| Ramachandran outliers (%) | 1.4 |
| B-factors: |  |
| Mean B-factor (protein atoms) (Å^2^) | 82.1 |
| Ligands (Å^2^) | 91.0 |
| PDB code | 9OTG |
